# Identification of the cellular transcription factor KLF16 as a novel repressive epigenetic repressor of HIV-1 transcription

**DOI:** 10.64898/2026.05.02.722432

**Authors:** Marion Santangelo, Maryam Bendoumou, Antoine Dutilleul, Soumia Khalfi, Estelle Plant, Christ Dominique Ngassaki-yoka, Lisa Pilosio, Caroline Vanhulle, Jonathan Dias, Tristan Marray, Antoine Fattaccioli, Marc Dieu, Jean-Pierre Routy, Olivier Rohr, Patricia Renard, Petronela Ancuta, Carine Van lint

**Author notes:** These authors contributed equally to this work (Marion Santangelo, Maryam Bendoumou, Antoine Dutilleul).

## Abstract

Despite antiretroviral therapy, human immunodeficiency virus type 1 (HIV-1) persists in latently-infected cells through epigenetic and transcriptional mechanisms. Latency-reversing agents have failed clinically, partly due to incomplete understanding of HIV-1 latency reversal. Here, using DNA-affinity capture and mass spectrometry on the HIV-1 5’ long terminal repeat (5’LTR) enhancer-core promoter, we identify KLF16 (Krüppel-like Factor 16) as a novel regulator of HIV-1 gene expression. KLF16 binds to the HIV-1 5’LTR *in vivo* at Sp1 binding sites, and KLF16 depletion reactivates latent HIV-1 in T-lymphoid and monocytic cell models. Mechanistically, KLF16 represses HIV-1 transcription by competing with Sp1 for promoter binding and by recruiting the Sin3A/HDAC1 and HP1α/Suv39H1 repressive epigenetic complexes. KLF16 is also upregulated in CD4^+^ T cells from ART-treated people with HIV-1 upon T-cell activation. Additionally, All-Trans retinoic acid (ATRA) reactivates latent HIV-1 in myeloid cells, partly by downregulating KLF16. These findings establish KLF16 as a novel transcriptional repressor of HIV-1, identifying it as a potential promising therapeutic target for cure strategies.

**Graphical abstract:** 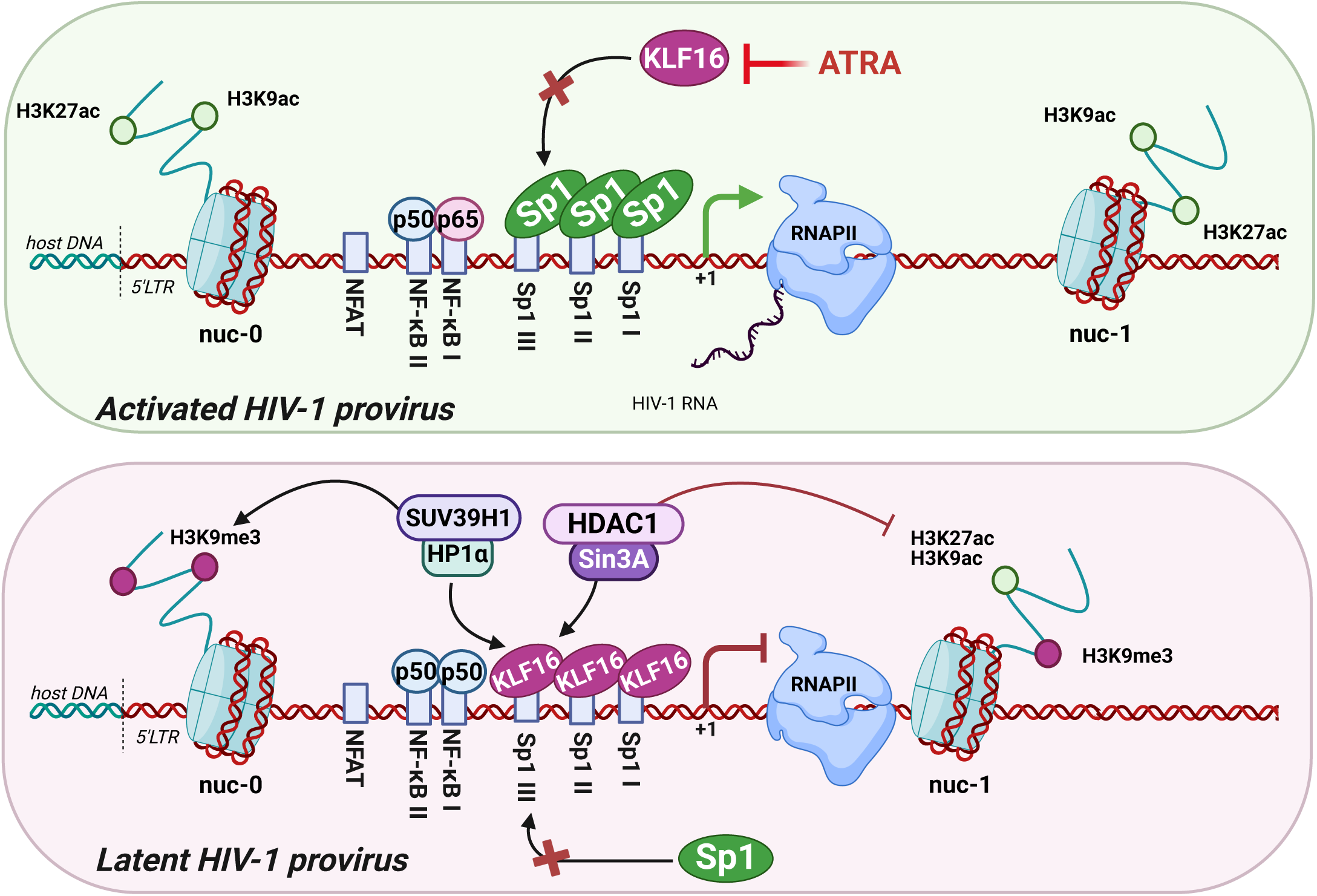

## Introduction

Despite effective antiretroviral therapy (ART), HIV-1 persists in cellular reservoirs composed of latently-infected cells that evade host immunity and can resume viral production upon treatment interruption^1–4^. The persistence of these reservoirs is linked to their heterogeneous nature, involving multiple cell types, such as long-lived resting memory CD4^+^ T cells and cells of myeloid origin^5–7^, and complex silencing mechanisms acting at the epigenetic, transcriptional, and post-transcriptional regulation levels^8–11^. To address HIV-1 latency, the “shock and kill” strategy proposes to use latency-reversing agents (LRAs) to reactivate HIV-1 gene expression and allow infected cells to die from viral cytopathic effects or immune-mediated killing (reviewed in^11–14^). However, while some LRAs have shown promise, most only reactivate HIV-1 expression without achieving measurable reservoir size reductions^15–17^, thereby highlighting the importance of still overlooked critical factors involved in HIV-1 gene expression, which is primarily regulated at the level of transcription.

HIV-1 transcription is a multifaceted process, governed by both the chromatin organization of the provirus and the array of *cis*-regulatory elements within the 5’-long terminal repeat (5’LTR)^17^. The interplay between chromatin accessibility and the binding of cellular transcription factors to this region ultimately determines whether HIV gene expression is enhanced or repressed. The core promoter and enhancer elements in the 5’LTR, which contain indispensable binding sites for transcription factors like NF-κB and Sp1^12,18^, are particularly critical hubs for this regulation. Indeed, Sp1 and NF-κB mediate recruitment to the 5’LTR of the cellular RNA polymerase II (RNAPII) transcription initiation complex. However, productive transcription is limited by promoter-proximal RNAPII pausing, which must be overcome by the HIV-1 protein Tat that recruits the positive transcription elongation factor b (P-TEFb) complex composed of the CDK9 kinase and Cyclin T1 to phosphorylate the RNAPII C-terminal domain (CTD) and negative elongation factors^19^. In order to identify novel cellular proteins that target the HIV-1 5’LTR enhancer and U3 core promoter region, we here conducted a proteomic screen of factors binding to a 100 bp probe centered on this region using a one-step DNA-affinity capture method combined with mass spectrometry. We identified a series of factors binding directly or indirectly to the DNA capture probe and selected KLF16 (Krüppel-like Factor 16) as the most promising candidate. KLF16 belongs to the family of Krüppel-like factors (KLFs) characterized by three highly conserved C2H2 zinc finger structures at their carboxy-terminus enabling their binding to GC-rich boxes, which are also recognized by Sp1^20,21^. KLF16 has been previously characterized as both a transcriptional repressor and activator, and KLF16-mediated transcriptional regulation is involved in many biological processes, such as cell growth, cell death, metabolism and cancer progression when overexpressed^22–29^. However, KLF16 functional role in HIV-1 transcriptional regulation has never been investigated so far. We showed that KLF16 promoted HIV-1 transcriptional silencing and latency by competing with Sp1 for binding to the 5’LTR and by recruiting epigenetic repressive complexes, positioning KLF16 as a novel actor in HIV-1 latency and a potential target for cure strategies.

## Results

### Identification of KLF16 as new potential HIV-1 regulator binding *in vitro* to the viral 5’LTR

To identify new cellular factors binding to the HIV-1 5’LTR and potentially able to regulate viral transcription, we conducted a screening using a one-step DNA-affinity capture method combined with an improved mass spectrometry process. We focused on both the enhancer and the U3 core promoter regions due to their important role in the recruitment of critical HIV-1 transcriptional regulators^12^. A 100 bp-long capture probe centered on these regions was incubated with nuclear extracts from either a T-lymphoid (Jurkat) or a myeloid (THP1) cell line (**Fig. 1a**). Data analysis allowed the identification of an initial pool of 692 proteins. This list was further refined using a custom R script designed to eliminate proteins with inconsistent detection across replicates, as well as technical contaminants and non-nuclear proteins. A final filter was applied to select for proteins whose DNA-binding consensus sequence was known, narrowing the list to nine high-confidence candidates. Using JASPAR analysis, these candidates were then mapped for potential binding sites within the DNA capture probe to identify novel regulators of HIV-1 transcription (**Fig. 1b**). Of these, only seven had predicted binding sites within the probe, including the known 5’LTR regulators JUND, NF-κB and TFAP4 (also known as AP-4) ^30^. We decided to focus our subsequent investigation on the transcription factor KLF16, because its potential binding sites were highly concentrated around the three critical Sp1 binding sites of the enhancer-core promoter region (**Fig. 1c**). These results are in agreement with the fact that KLF16 belongs to the Sp1-like protein family and has a consensus sequence similar to that of the Sp1 protein^21^. KLF16 has been characterized as both a transcriptional repressor and activator of several cellular promoters^26,31^ and its role in HIV-1 transcriptional regulation had never been investigated so far.

**Figure 1.**
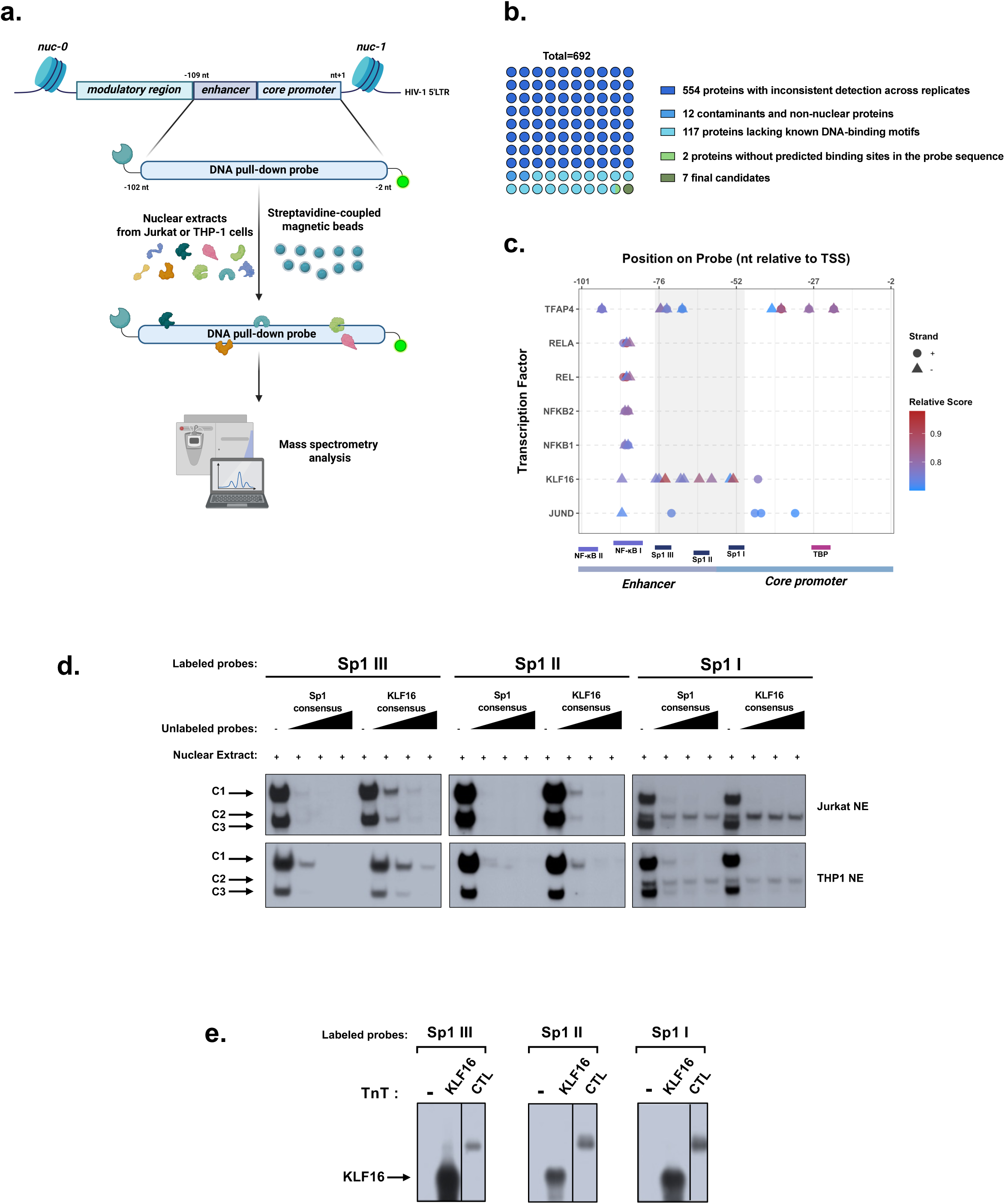
Identification of KLF16 as new potential HIV-1 regulator binding in vitro to the HIV-1 5’LTR. **(a)** DNA pull-down assays were performed by incubating a specific biotinylated probe spanning both the enhancer region and the U3 core promoter of the HIV-1 5’LTR with 1 mg of either Jurkat or THP-1 nuclear extracts. Streptavidin-coupled magnetic beads enabled purification of proteins binding to the DNA capture probe. Proteins were analysed by mass spectrometry. **(b)** Sequence coverage shown for KLF16. **(c)** Pie chart representing the systematic filtering strategy based on detection consistency, absence of known contaminants, and presence of DNA-binding motifs to identify candidate transcription factors binding to the DNA pull-down probe. **(d)** Map showing the localization of predicted binding sites for the seven candidate transcription factors (identified in **c**) within the HIV-1 54LTR enhancer-core promoter region relative to the transcription start site (TSS = nt +1). Dots and triangles represent binding sites on the positive and negative DNA strands, respectively. Colours indicate predicted binding scores from JASPAR analysis (red = highest affinity; blue = lowest affinity). **(e)** Radiolabeled probes corresponding to the 1st (Sp1 I), 2nd (Sp1 II), and 3rd (Sp1 III) Sp1 binding sites of the HIV-1 5’LTR U3 region were incubated with 10 μg of either Jurkat (top panel) or THP-1 (bottom panel) nuclear extracts (NE) in the absence of competitor or in the presence of increasing molar excesses of unlabeled consensus sequences for either Sp1 or KLF16. **(f)** The radiolabelled probes corresponding to the 1st, 2nd, or 3rd Sp1 binding site sequences were incubated with KLF16 pure extract or luciferase protein as negative control. One representative experiment out of three independent replicates is shown. Only the specific retarded bands of interest are displayed.

Further *in silico* analyses using the Jaspar algorithm^32,33^ suggested that the three Sp1 binding sites of the HIV-1 U3 core promoter could correspond to three KLF16 binding sites. Therefore, we next performed electrophoretic mobility shift assays (EMSAs) using three probes corresponding to the three Sp1 binding sites of the HIV-1 U3 core promoter (termed Sp1 I (nt -55 to nt -46 where nt +1 corresponds to the TSS), Sp1 II (nt -65 to nt -57), and Sp1 III (nt -76 to nt -67), respectively). These radiolabeled probes were incubated with nuclear extracts prepared from either CD4+ T-lymphocytic (Jurkat) or monocytic (THP1) cell lines, in absence or in presence of increasing molar excesses of unlabeled double-stranded oligonucleotides, serving as competitors and corresponding either to the Sp1 binding site consensus sequence (termed Sp1 consensus) or to the KLF16 binding site consensus sequence (termed KLF16 consensus) (**Fig. 1d**). Incubation of the radiolabeled probes with the nuclear extracts resulted in the formation of two retarded complexes for the 3^rd^ (Sp1 III) and 2^nd^ Sp1 (Sp1 II) site probes and three complexes for the 1^st^ Sp1 (Sp1 I) site probe with either Jurkat or THP1 nuclear extracts (**Fig. 1d**). Except for the second retarded complex observed with the 1^st^ Sp1 site probe, the formation of the other retarded complexes was inhibited by competition with increasing molar excesses of both unlabeled Sp1 and KLF16 consensus oligonucleotides, demonstrating the specificity of these DNA-protein complexes to the Sp1 and KLF16 consensus motif **(Fig. 1d).** Taken together, our *in vitro* binding studies demonstrated that proteins present in the retarded complexes were able to bind to both the Sp1 and KLF16 consensus motifs and could include KLF16.

To investigate the direct *in vitro* binding of KLF16 to these three Sp1 binding sites, we performed EMSAs with the three Sp1 radiolabeled probes incubated with *in vitro* produced KLF16 or luciferase as a negative control (termed KLF16 or CTL-, respectively). This resulted in the formation of one specific retarded complex designated KLF16 compared to the negative control **(Fig. 1e).**

Taken together, we demonstrated for the first time that KLF16 directly binds *in vitro* to the three consecutive Sp1 binding sites located within the U3 region core promoter of the HIV-1 5’LTR.

### KLF16 is recruited *in vivo* to the HIV-1 5’LTR and represses viral gene expression

To investigate potential cell type specific mechanisms of HIV latency, we developed parallel cellular models for HIV-1 latency, as no simultaneously generated latent cell lines of both T-lymphoid and myeloid origins infected with the same viral strain were available. We infected Jurkat and THP-1 cells with the dual-reporter virus HIVGKO, which allows for the discrimination of latently-infected cells (csGFP⁻/mKO2⁺) from productively infected cells (csGFP⁺/mKO2⁺)^34^. Clonal populations of these latently-infected cells were then isolated via fluorescence-activated cell sorting (FACS) and expanded to establish stable cell lines, termed Jurkat HIVGKO and THP HIVGKO. Using these cell lines, we investigated the *in vivo* recruitment of KLF16 to the HIV-1 5’LTR by chromatin immunoprecipitation (ChIP) assays using primers hybridizing specifically to the HIV-1 5’LTR, in the Nuc-1 region. As shown in **Fig.2a**, KLF16 was recruited to the latent promoter in both Jurkat HIVGKO and THP HIVGKO cell lines. Furthermore, we observed similar results in primary CD4^+^ T cell and monocyte-derived-macrophage (MdM) models for HIV-1 infection, supporting the hypothesis that KLF16 could be involved in HIV-1 transcriptional regulation (**Fig. 2a**).

**Figure 2.**
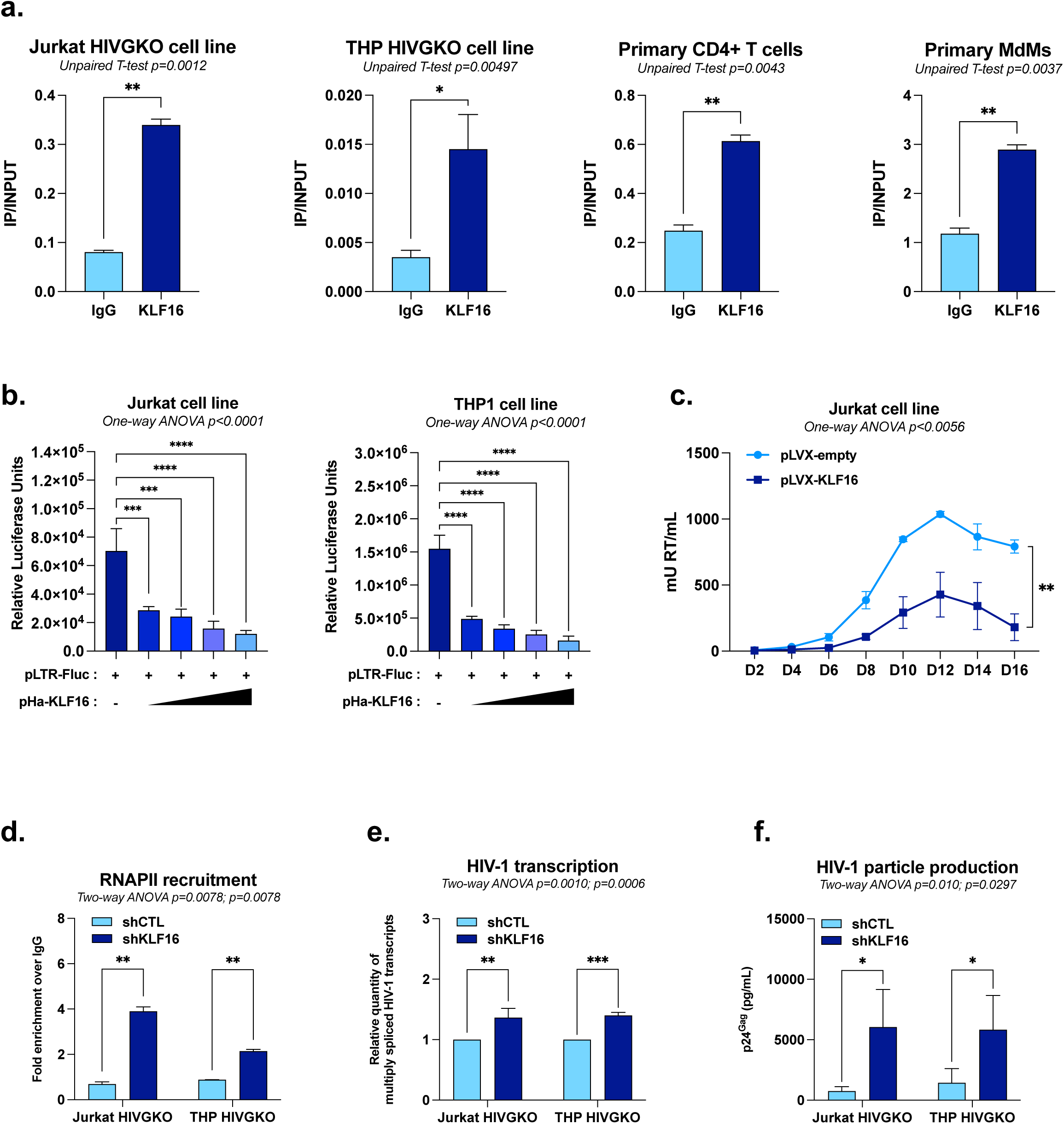
KLF16 is recruited in vivo to the HIV-1 5’LTR and represses viral gene expression. **(a)** Chromatin samples were prepared from Jurkat HIVGKO and THP HIVGKO cell lines or from primary CD4^+^ T cells and monocyte-derived macrophages (MDMs) infected with HIV-1 and immunoprecipitated with a specific antibody against KLF16 or with a non-specific IgG serving as background control. Results are presented as histograms indicating the ratio of IP/INPUT. **(b)** Jurkat or THP-1 cells were transiently transfected with p-LTR-Fluc and increasing doses of an expression vector for KLF16 (pHA-KLF16) or equimolar amounts of the empty parental vector pcDNA3.1. Luciferase activities were measured and normalized by protein concentration. **(c)** Uninfected Jurkat cell lines stably overexpressing either the KLF16 expression vector (pLVX-KLF16) or the control vector (pLVX-empty) were infected with a fully replicative HIV-1 molecular clone, and viral particle production was monitored over time by quantifying RT activity in culture supernatants using SG-PERT assay. Jurkat HIVGKO and THP HIVGKO cell lines were transduced with either shCTL or shKLF16 and were used for **(d)** chromatin immunoprecipitation assays with a specific antibody against RNAPII or a non-specific IgG serving as background control (values represent fold enrichments over IgG) and **(e)** total RNA extraction to quantify HIV-1 multiply-spliced (MS) mRNA levels by RT-qPCR. **(f)** Culture supernatants from either shCTL- or shKLF16-transduced Jurkat HIVGKO and THP HIVGKO cells were analysed for viral production by measuring p24 capsid protein by ELISA. One representative experiment out of three independent replicates is shown. Statistical analysis methods and corresponding *p* values are indicated above each graph.

To investigate the functional role of KLF16 in HIV-1 transcription, we subcloned the HIV-1 5’LTR region in an episomal reporter construct where the 5’LTR controls the firefly luciferase gene (referred to as pLTR-Fluc)^35^. Jurkat and THP1 were transiently co-transfected with the reporter construct along with increasing doses of a KLF16 expression vector (referred to as pHA-KLF16). Overexpression of KLF16 led to a dose-dependent and statistically significant decrease in the relative luciferase activity compared to the control condition in both T-lymphocytic and monocytic cellular contexts (**Fig. 2b**). Because HIV-1 transcription is highly sensitive to RNAPII promoter-proximal pausing and requires the presence of the viral protein Tat which recruits the P-TEFb complex to promote productive transcriptional elongation, we next assessed whether KLF16 also modulates this Tat-activated state. To this end, HEK293T cells were transiently co-transfected with the LTR reporter construct, a Tat expression vector, and increasing doses of the KLF16 expression vector (pHA-KLF16). Overexpression of HA-KLF16 induced a statistically significant decrease in Tat-mediated trans-activation of HIV-1 5’LTR-driven gene expression (**Extended data Fig. 1a**). These data showed that KLF16 represses HIV-1 5’LTR promoter activity in presence and in absence of Tat.

To further evaluate the potential role of KLF16 in the establishment of HIV-1 latency, we generated two Jurkat-derived cell lines transduced with either a lentiviral vector encoding for KLF16 (pLVX-KLF16) or the corresponding empty control vector (pLVX-empty) and selected for puromycin resistance. After confirming KLF16 overexpression by immunoblot (**Extended Data Fig. 1b**), we infected the cell lines with a replication-competent HIV-1 molecular clone (pNL4.3-GFP) and HIV replication kinetics was monitored over 16 days by measuring viral particle production *via* SG-PERT assays^36^. Our results demonstrated that constitutive overexpression of KLF16 reduces in a statistically significant manner the production of HIV-1 particles starting at day 8 after infection (**Fig. 2c**). We further quantified HIV-1 integrated DNA at day two post-infection and observed no significant difference between the two conditions. This indicates that the KLF16-mediated reduction in viral replication does not result from altered proviral integration and likely stems from an effect on HIV-1 transcription (**Extended Data Fig. 1c**).

To determine the role of KLF16 in establishing or maintaining HIV-1 transcriptional silencing at the viral promoter during latency, we downregulated endogenous KLF16 using RNA interference. Latently-infected Jurkat and THP HIVGKO cells were stably transduced with lentiviral vectors expressing either an shRNA targeting KLF16 mRNA (shKLF16) or a non-targeting control shRNA (shCTL). After validating KLF16 knockdown in puromycin-selected clones by RT-qPCR (**Extended Data Fig. 1d**), we observed several indicators of increased viral activity. First, ChIP experiments revealed that RNAPII recruitment to the 5’LTR was significantly increased in KLF16-downregulated cells (**Fig. 2d**). Second, consistent with this, RT-qPCR quantification demonstrated a statistically significant increase in the levels of multiply spliced (elongated) HIV-1 transcripts (**Fig. 2e**). Third, quantification of the p24^Gag^ capsid protein by ELISA demonstrated that KLF16 knockdown results in increased HIV-1 production in culture supernatants (**Fig. 2f**).

Finally, since *cis*-regulatory regions beyond the 5’LTR, including the *pol* gene intragenic *cis*-regulatory region (IRR) and the 3’LTR, contain Sp1 binding sites that recruit Sp1 and regulate HIV-1 transcription^30^, we evaluated whether KLF16 could also bind *in vivo* to these regions *via* recognition of Sp1 motifs using ChIP-qPCR in primary CD4^+^ T cells and MdMs infected with HIV-1. KLF16 was recruited at the 5’LTR, but not at the IRR or 3’LTR, in primary CD4^+^ T cells (**Extended Data Fig. 1e**). Similarly, in MdMs, KLF16 bound robustly to the 5’LTR, with only slight recruitment at the IRR and none at the 3’LTR (**Extended Data Fig. 1f**). Given that other retroviruses including Human T-Lymphotropic Virus 1 (HTLV-1) and endogenous retroviruses such as HERV-K also contain Sp1 binding sites in their promoter and rely on Sp1 for transcriptional regulation^37,38^ , we assessed KLF16 recruitment to these Sp1 sites but found no binding to either HTLV-1 or HERV-K 5’LTRs in the HTLV-1 latently-infected cell line TLOM-1 or in primary CD4^+^ T cells and MdMs, respectively (**Extended Data Fig. 1e-g**). These results demonstrate that KLF16 specifically targets the HIV-1 5’LTR promoter *via* Sp1-motif recognition to repress viral transcription.

Taken together, our results demonstrate that KLF16 is involved in the establishment and maintenance of HIV-1 latency, and that its functional role takes place, at least in part, at the transcriptional level.

### KLF16 competes with Sp1 for HIV-1 5’LTR binding to repress viral gene expression

Since KLF16 binds *in vitro* to the three Sp1 binding sites within the HIV-1 U3 core promoter (**Fig. 1d**), we investigated a potential competitive binding mechanism between the two transcription factors. First, we showed the importance of these sites by introducing mutations in all known Sp1 binding motifs within the 5’LTR region into two distinct 5’LTR-luciferase reporter constructs: one mutated in the three Sp1 sites of the U3 region (referred to as pLTR-FlucΔSp1) and the second mutated in the two Sp1 sites of the leader region (referred to as pLTR-FlucΔSp1 Leader). Consistent with the literature^8,39^, mutations in the three Sp1 binding sites of the HIV-1 U3 region, but not of those in the leader region, affected basal 5’LTR promoter activity (**Extended Data Fig. 2a**). The KLF16-mediated repression of LTR activity was abolished when the three U3 core promoter Sp1 sites were mutated, but not when the two Sp1 sites of the leader region were mutated, thereby highlighting the critical role of the promoter-proximal Sp1 sites for KLF16 function (**Fig. 3a**).

**Figure 3.**
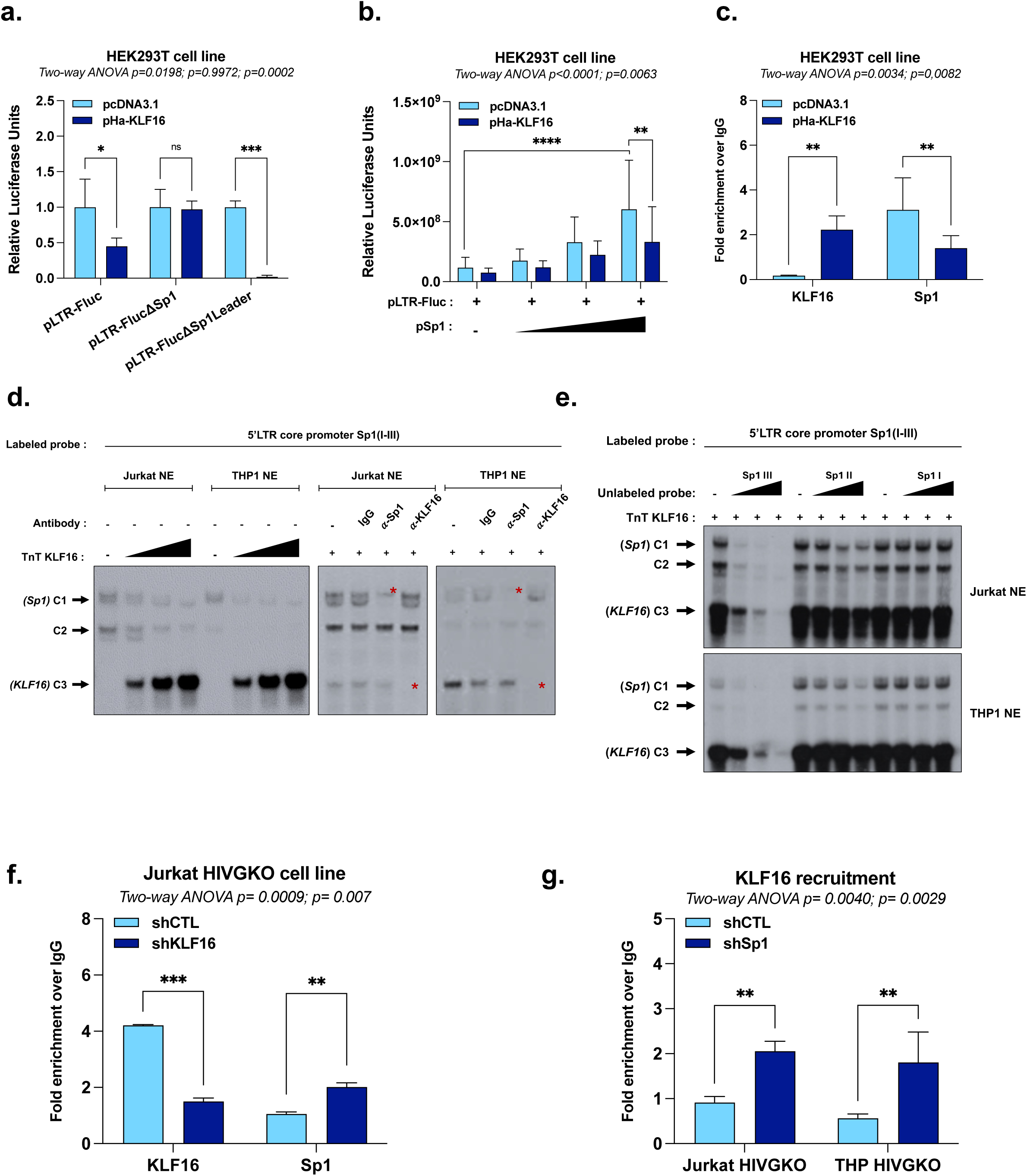
KLF16 competes with Sp1 for HIV-1 5’LTR binding to repress HIV-1 gene expression. **(a)** HEK293T cells were transiently transfected with either p-LTR-Fluc, pLTR-FlucΔSp1, or pLTR-FlucΔSp1Leader together with a single dose of pHA-KLF16 or the empty control vector pcDNA3.1. Luciferase activities were measured and normalized by protein concentrations. **(b)** HEK293T cells were transiently transfected with p-LTR-Fluc and increasing doses of the Sp1 expression vector (pSp1) in the presence of either a single dose of pHA-KLF16 or the empty control vector pcDNA3.1. Luciferase activities were measured and normalized by protein concentrations. **(c)** Chromatin samples prepared from the experiment in (b) (HEK293T cells transfected with p-LTR-Fluc and the highest dose of pSp1 alone or together with a single dose of pHA-KLF16) were immunoprecipitated with specific antibodies against KLF16, Sp1, or with a non-specific IgG serving as background control (values represent fold enrichments over IgG). **(d)** Radiolabeled probe spanning the 1st (Sp1 I), 2nd (Sp1 II), and 3rd (Sp1 III) Sp1 binding sites of the HIV-1 5’LTR U3 region was incubated with 10 μg of either Jurkat or THP-1 nuclear extracts (NE) in the presence or absence of increasing doses of recombinant KLF16 protein. Supershifted complexes are indicated by a red asterisk. **(e)** Radiolabeled probe spanning the 1st (Sp1 I), 2nd (Sp1 II), and 3rd (Sp1 III) Sp1 binding sites of the HIV-1 5’LTR U3 region was incubated with 10 μg of either Jurkat or THP-1 nuclear extracts and a single dose of recombinant KLF16 protein in the absence of competitor or in the presence of increasing molar excesses of unlabelled consensus sequences corresponding to either the 1^st^, the 2^nd^, or the 3^rd^ Sp1 binding site. **(f)** Jurkat HIVGKO cells transduced with either shCTL or shKLF16 were used for chromatin immunoprecipitation assays with specific antibodies against KLF16, Sp1, or a non-specific IgG serving as background control (values represent fold enrichments over IgG). **(g)** Jurkat HIVGKO and THP HIVGKO cells transduced with either shCTL or shSp1 were used for chromatin immunoprecipitations with a specific antibody against KLF16 or with a non-specific IgG serving as background control (values represent fold enrichments over IgG). One representative experiment out of three independent replicates is shown. Statistical analysis methods and corresponding *p* values are indicated above each graph.

To functionally evaluate the competition, we co-transfected HEK293T cells with the pLTR-Fluc reporter construct and increasing doses of a Sp1 expression vector (referred to as pSp1), with or without a fixed dose of the KLF16 expression vector. While Sp1 overexpression alone enhanced LTR promoter activity, the co-expression of KLF16 significantly antagonized this effect, reducing Sp1-mediated activation by approximately 50% (**Fig. 3b**). ChIP-qPCR experiments further demonstrated this competitive interaction since KLF16 overexpression increases its own recruitment to the 5’LTR while concurrently decreasing Sp1 recruitment (**Fig. 3c**).

To further characterize the competition between KLF16 and Sp1 at the DNA-protein interface, we performed EMSAs. A radiolabeled probe spanning the three Sp1 binding sites of the HIV-1 U3 core promoter was incubated with nuclear extracts from either Jurkat or THP1 cell lines, resulting in the formation of two retarded protein-DNA complexes (C1 and C2). The addition of *in vitro* produced KLF16 protein caused a dose-dependent decrease in the intensity of these complexes and the concomitant formation of a new, slower-migrating complex (C3). Supershift analysis showed that complex C1 contained Sp1, while complex C3 contained KLF16, directly demonstrating that KLF16 displaced a Sp1-containing complex from its binding sites (**Fig. 3d**). To map the specific sites of this competition, we next performed EMSAs using as competitors unlabeled oligonucleotide probes corresponding to each of the three individual Sp1 binding sites. Only molar excesses of unlabeled probes for the 3^rd^ and, to a lesser extent, the 2^nd^ Sp1 site were able to compete away the retarded complexes. In contrast, the probe for the 1^st^ Sp1 site had no effect (**Fig. 3e**). Together, these results indicate that KLF16 directly competes with Sp1 for binding to the HIV-1 promoter, with a clear preference for the first two Sp1 binding sites within the U3 region.

To functionally assess the competitive binding between KLF16 and Sp1 under endogenous conditions, we performed reciprocal knockdown experiments coupled with ChIP-qPCR analysis in our cell line models for HIV-1 latency. First, shRNA-mediated downregulation of KLF16 in Jurkat HIVGKO cells resulted in significantly increased recruitment of Sp1 to the 5’LTR (**Fig. 3f**). Conversely, downregulation of Sp1 in both Jurkat and THP HIVGKO cell lines, validated by RT-qPCR (**Extended Data Fig. 2b**), led to a concomitant increase in KLF16 recruitment to the viral promoter (**Fig. 3g**). Taken together, these results demonstrate that KLF16 and Sp1 compete for binding at the HIV-1 promoter *in vivo*.

### KLF16 modulates the epigenetic profile of the HIV-1 5’LTR to repress viral gene expression

The epigenetic state of the HIV-1 5′LTR is a critical determinant of latency, where histone hypermethylation and hypoacetylation cause chromatin compaction by recruiting histone deacetylases (HDAC) and methyltransferases (HMT), blocking access to transcription machinery and maintaining viral latency^13^. Because KLF16 had been previously reported to recruit the repressive Sin3A-HDAC1 complex to cellular promoters^29^, we hypothesized that it could similarly regulate HIV-1 gene expression by modulating the epigenetic profile of the 5’LTR. To test this hypothesis, we used shRNA to downregulate KLF16 expression in our latently-infected Jurkat and THP HIVGKO cell lines, as described before. ChIP-qPCR assays revealed that the KLF16 knockdown, which induced HIV-1 reactivation (**Fig. 2d-f**), was associated with a significant decrease in both Sin3A and HDAC1 recruitment to the 5’LTR in both cellular contexts (**Fig. 4a and 4c**). Accordingly, this loss of epigenetic repressive machinery was accompanied by a corresponding increase in the activating histone marks H3K9ac and H3K27ac at the viral promoter (**Fig. 4b and 4d**). RT-qPCR analysis indicated that these effects were due to changes in protein recruitment, as the mRNA levels of Sin3A and HDAC1 themselves were unaffected by KLF16 downregulation (**Extended Data Fig. 3a-b**).

**Figure 4.**
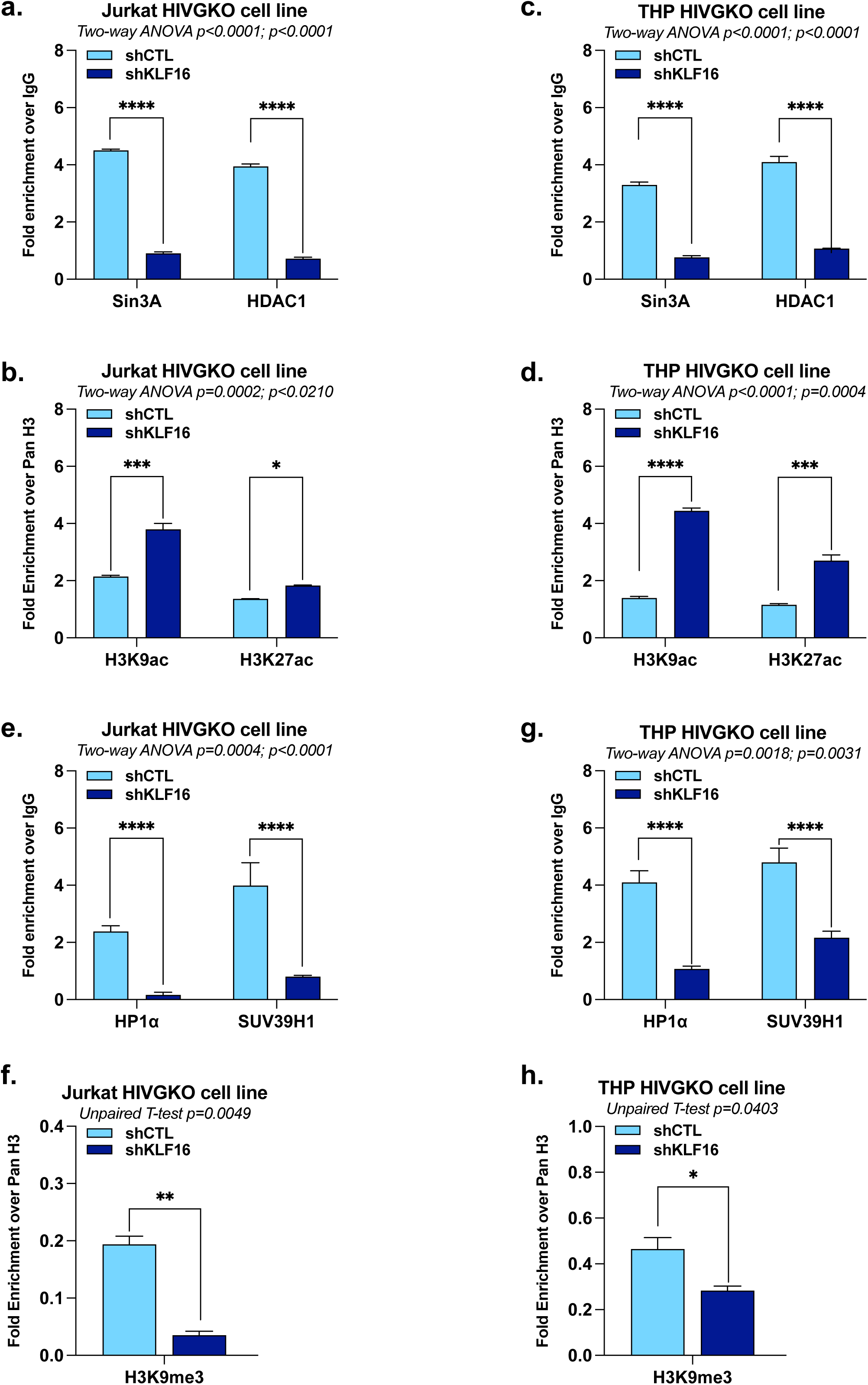
KLF16 modulates the epigenetic profile of the HIV-1 5’LTR to repress viral gene expression. Jurkat HIVGKO and THP HIVGKO cells transduced with either shCTL or shKLF16 were used for chromatin immunoprecipitations with a non-specific IgG serving as background control or with **(a)-(c)** specific antibodies targeting Sin3A or HDAC1, or **(e)-(g)** specific antibodies targeting HP1α or SUV39H1 (values represent fold enrichments over IgG). The same chromatin samples were used to assess the histone modification profile at the HIV-1 5’LTR using specific antibodies targeting total histone H3 (Pan H3) or **(b)-(d)** H3K9ac and H3K27ac, or **(f)-(h)** H3K9me3 (values represent fold enrichments over Pan H3). One representative experiment out of three independent replicates is shown. Statistical analysis methods and corresponding *p* values are indicated above each graph.

Additionally, given the mechanistic similarities KLF16 shares with other Krüppel-like factors such as KLF11^40^, which directly recruits the HP1α-SUV39H1 complex to promoters, leading to H3K9 trimethylation and transcriptional repression^41^, we investigated whether KLF16 also modulates the methylation status of the HIV-1 promoter. We performed ChIP-qPCR in Jurkat and THP HIVGKO cells following shRNA-mediated downregulation of KLF16. Our results showed that KLF16 depletion is associated with a decreased recruitment of both HP1α and SUV39H1 to the 5’LTR (**Fig. 4e and 4g**). This is accompanied by a significant decrease in the level of the repressive histone mark H3K9me3 at the HIV-1 promoter (**Fig. 4f and 4h**). RT-qPCR analysis showed that these effects are due to changes in protein recruitment, as the mRNA levels of HP1α and SUV39H1 themselves were unaffected by KLF16 downregulation (**Extended Data Fig. 3a-b**).

Taken together, our data demonstrate that KLF16 mediates the *in vivo* recruitment of two distinct epigenetic silencing complexes, Sin3A-HDAC1 and HP1α-SUV39H1, to the HIV-1 promoter to establish a local repressive chromatin environment. This highlights a bimodal mechanism for KLF16 in silencing HIV-1 transcription, acting both through repressive epigenetic modifications and through its competition with Sp1 for promoter binding.

### KLF16 represses HIV-1 gene expression in primary memory CD4^+^ T cells infected *in vitro*

While latently-infected cell lines serve as valuable model systems for mechanistic studies, they may not fully capture the spectrum and heterogeneity of HIV-1 latency mechanisms occurring *in vivo*. To explore the function of KLF16 in a more physiologically relevant context, we evaluated its role in a primary CD4^+^ T cell model of HIV-1 infection. Memory CD4^+^ T cells isolated from HIV-uninfected donors (People without HIV (PWOH)) were activated, nucleofected with either a siRNA targeting KLF16 (siKLF16) or a non-targeting control (siNT), and subsequently single-round infected with a VSV-G pseudotyped HIV-1 strain (**Fig. 5a**). After validating efficient KLF16 downregulation by RT-qPCR (**Fig. 5b**), we observed that the levels of integrated HIV-1 DNA were significantly reduced in siKLF16-treated cells compared to control cells, suggesting a potential role for KLF16 in pre-integration processes (**Fig. 5c**). To discriminate the effects on pre-integration from those on post-integration, we quantified extracellular viral particle (p24^Gag^) production and normalized these values to the amount of integrated HIV-DNA. This analysis revealed that, on a per-provirus basis, KLF16 depletion led to a significant increase in HIV particle production (**Fig. 5d**). These findings demonstrate that KLF16 functions as a repressor of HIV-1 gene expression in infected primary CD4^+^ T cells, independently of its potential effects on the pre-integration steps of viral replication cycle.

**Figure 5.**
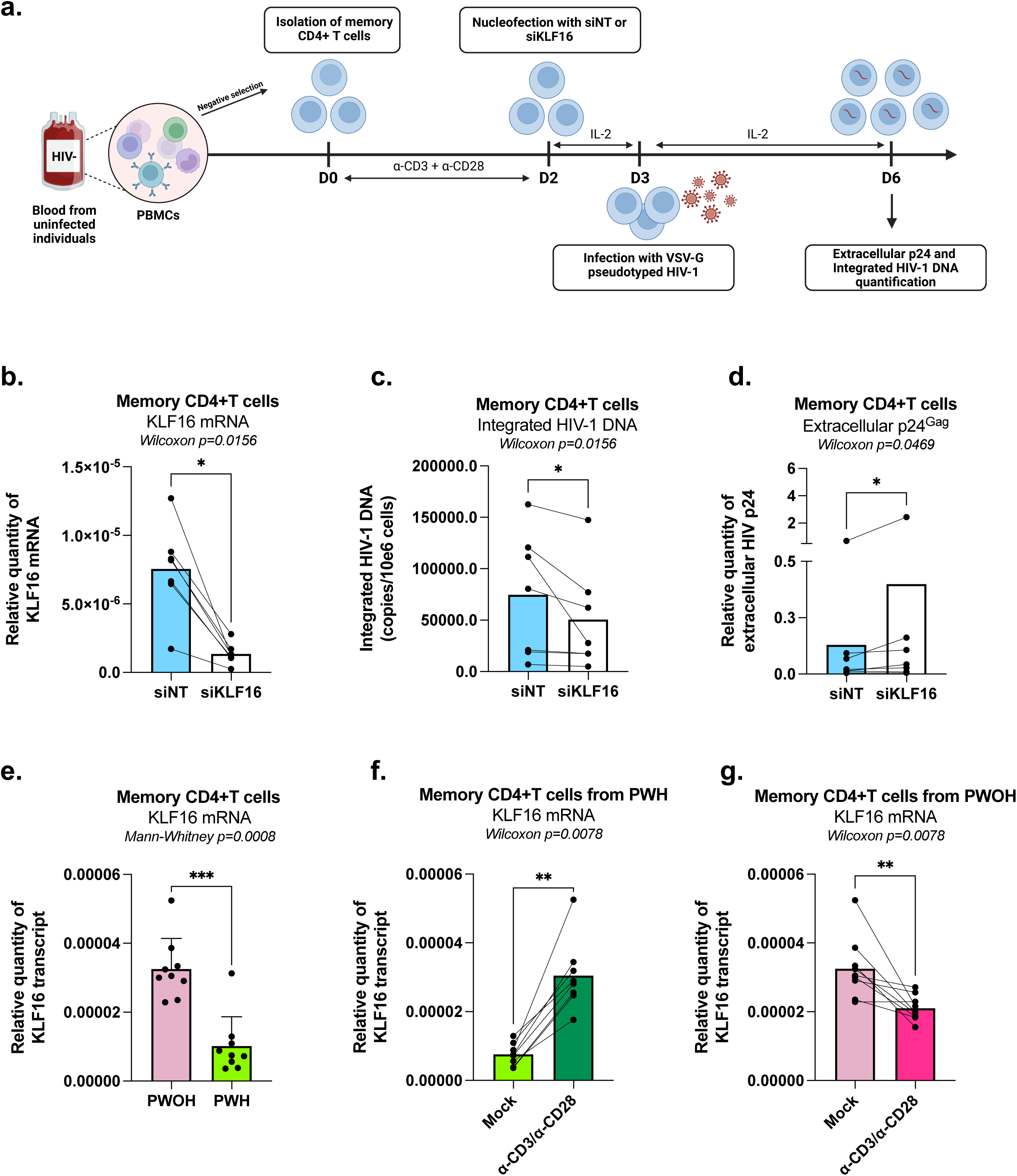
KLF16 represses HIV-1 gene expression in primary CD4+ T cells infected *in vitro* and KLF16 expression is upregulated following TCR-triggering in memory CD4+ T cells from PWH. **(a)** Experimental flowchart. Memory CD4^+^ T cells were isolated from PBMCs of PWOH were stimulated with α-CD3+α-CD28 antibodies for 3 days and then were nucleofected with non-targeting control (siNT) or KLF16-targeting (siKLF16) small interfering RNA (siRNA). Nucleofected cells were cultured in the presence of rhIL-2 (5 ng/ml) for 24 hours, then infected with VSV-G-pseudotyped HIV-1 encoding GFP (50 ng HIV-p24/10e6 cells) and cultured in the presence of rhIL-2 for an additional 3 days. **(b)** KLF16 mRNA expression was measured by RT-qPCR 24 hours post-nucleofection. **(c)** At day 3 post-infection, cells were harvested to quantify the levels of integrated HIV DNA by nested real-time PCR. **(d)** Levels of HIV-p24 capsid protein were measured by ELISA in cell culture supernatants collected at day 3 post-infection. Results were generated with cells from n=7 PWOH participants. **(e)- (g)** Memory CD4^+^ T cells isolated from PBMCs of PWOH (n = 9) or of ART-treated PWH (n = 9) were stimulated with α-CD3+α-CD28 antibodies or left unstimulated for 18 hours and used for the measurement of KLF16 mRNA expression. **(e)** KLF16 mRNA levels quantified by RT-qPCR in samples from PWOH and PWH *ex vivo*. **(f)** KLF16 mRNA levels quantified by RT-qPCR in samples from PWH upon TCR triggering *in vitro*. **(g)** KLF16 mRNA levels quantified by RT-qPCR in samples from PWOH upon TCR triggering *in vitro*. Statistical analysis methods and corresponding *p* values are indicated above each graph.

### KLF16 expression is promoted following TCR activation of memory CD4^+^ T cells from ART-treated PWH

Given our observation that KLF16 acts as a repressor of HIV-1 gene expression and that its depletion promotes proviral reactivation, we sought to examine whether its expression was altered in the context of chronic HIV infection. To this end, KLF16 expression in memory CD4^+^ T cells isolated from PWOH and from ART-treated PWH was compared to assess whether T cell receptor (TCR)-triggering could modulate KLF16 expression in infected cells. KLF16 expression was markedly lower in CD4^+^ T cells from ART-treated PWH compared to PWOH *ex vivo* (**Fig. 5e**, 2.87-fold decrease). A decrease in KLF16 expression may be likel^42^y due to persistent immune activation and transcriptional reprogramming that are known to occur in CD4^+^ T cells from PWH during ART^43,44^. Interestingly, upon polyclonal TCR stimulation using anti-CD3+anti-CD28 antibodies, KLF16 expression in memory CD4^+^ T cells from PWH increased to levels similar to those observed in memory CD4^+^ T cells from PWOH **(Fig. 5f**, 2.81-fold increase). These findings suggest that, while baseline KLF16 levels maybe insufficient to maintain robust HIV-1 latency in PWH, TCR-mediated upregulation of KLF16 in memory CD4^+^ T cells may help controlling the magnitude of viral transcription and limiting viral outgrowth from reservoir cells. In contrast, in memory CD4^+^ T cells from PWOH, TCR triggering reduced the levels of KLF16 transcripts (**Fig. 5g**, 1.54-fold decrease), consistent with the documented effect of TCR-triggering on the ability of T cells to support productive HIV-1 infection^42^.

Altogether, these results suggest that HIV-1 infection and/or long-term ART regimens may select for memory CD4^+^ T cells capable of upregulating KLF16 expression following activation, thereby dampening the HIV-1 reactivation potential and contributing to viral reservoir persistence.

### Retinoic acid, a modulator of gut-homing, activates HIV-1 transcription in myeloid cells in part through KLF16 downregulation

The gut represents a major reservoir for HIV-1 persistence^45,46^, where retinoic acid (RA) produced by intestinal dendritic cells promotes both lymphocyte gut-homing and heightened HIV-1 transcriptional activity in myeloid cells^1–5^. Therefore, elucidating the molecular mechanisms underlying RA-induced HIV transcriptional activation could reveal novel therapeutic targets to reduce this persistent reservoir. Our prior RNA-sequencing experiments in primary MdMs identified KLF16 as a gene downregulated following ATRA treatment (see the GEO database under accession number GSE226653)^47^, suggesting that ATRA may activate HIV-1 expression, in part, through KLF16 downregulation. We validated this effect by RT-qPCR in primary MdMs exposed to ATRA (**Fig. 6a**). We further investigated this effect by treating our HIV-1 latently-infected T-lymphocytic and monocytic cell lines with ATRA and we observed that KLF16 mRNA was significantly reduced only in monocytic cells (**Fig. 6b**), whereas HIV-1 transcription and replication were increased in both cell types (**Fig.6c-d**). These findings imply that ATRA-mediated HIV-1 reactivation can partially occur *via* inhibition of KLF16 in myeloid cells. Accordingly, epigenetic profiling at the HIV-1 5’LTR revealed that ATRA treatment increased activating histone marks (H3K9ac, H3K27ac) and decreased the repressive mark H3K9me3 (**Fig. 6e**).

**Figure 6.**
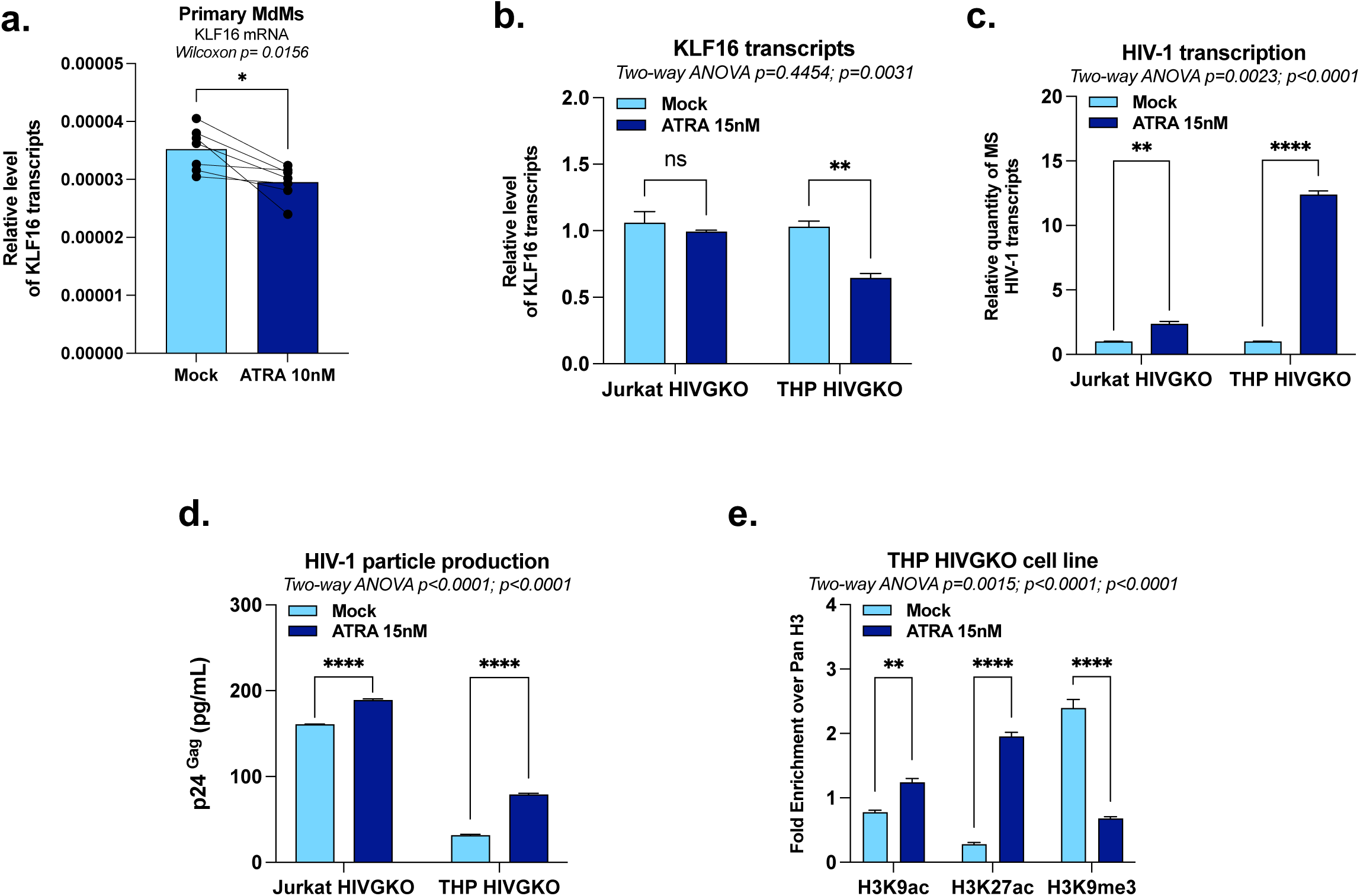
ATRA activates HIV-1 gene expression in myeloid cells in part through KLF16 downregulation. **(a)** Monocyte-derived macrophages (MDMs) were obtained by culturing monocytes isolated from PWOH in medium containing M-CSF (20 ng/mL) for 6 days and then treated or not with 10 nM ATRA. Total RNA was extracted to quantify KLF16 mRNA levels by RT-qPCR (n=7 PWOH). Jurkat HIVGKO and THP HIVGKO cells were treated with 15 nM of ATRA for 24 hours, and **(b)-(c)** a fraction of cells was used for total RNA extraction to quantify KLF16 mRNA and HIV-1 MS mRNA levels by RT-qPCR, and **(d)** culture supernatants were analysed for viral production by measuring p24 capsid protein by ELISA. **(e)** The remaining cell fraction was used for chromatin preparation to evaluate the histone modification profile at the HIV-1 5’LTR using antibodies targeting total histone H3, H3K9ac, H3K27ac, or H3K9me3. One representative experiment out of three independent replicates is shown. Statistical analysis methods and corresponding *p* values are indicated above each graph.

Altogether, our results demonstrate that ATRA-mediated HIV-1 reactivation coincides with KLF16 downregulation in primary MdMs and THP HIVGKO cells and modulates the epigenetic landscape of the viral promoter, supporting a role for KLF16 in the reversal of HIV-1 latency in myeloid cells.

## Discussion

Although potent LRAs can reactivate HIV-1 gene expression *in vitro*, they rarely reduce the reservoir size *in vivo*, reflecting the heterogeneous nature of HIV-1 latency^48^, and emphasizing the need to better understand the full range of factors promoting viral latency. In the present study, we identified the host transcriptional regulator KLF16 as a novel epigenetic repressor of HIV-1 transcription in both T-lymphocytic and monocytic latently-infected cells. Thus, KLF16 could constitute a novel molecular target for HIV-1 cure strategies.

Functionally, we showed that KLF16 repressed HIV-1 transcription both through competition with Sp1 for binding to the three critical Sp1 binding motifs of the viral 5’LTR core promoter and through recruitment of the repressive epigenetic complexes Sin3A-HDAC1 and HP1a-SUV39H1 to promote heterochromatin formation at the HIV-1 promoter. Moreover, the repressive activity required exclusively the three Sp1 binding sites located in the 5’LTR U3 core promoter, which are indispensable for both activation and silencing of HIV-1 transcription^18,30^. During productive infection, Sp1 binds to these GC-rich *cis*-regulatory elements and cooperates with adjacent NF-κB to recruit p300/CBP, P-TEFb, Mediator, and the basal transcription machinery, thereby promoting histone acetylation, relieving promoter-proximal pausing of RNA polymerase II, and driving robust transcription elongation^49–52^. By contrast, these same Sp1 sites also mediate transcriptional repression through two main mechanisms. First, Sp1 can recruit co-repressor proteins: in CD4⁺ T cells, Sp1 associates with c-Myc to bring in HDAC1/2^53^, while in microglial cells, Sp1 partners with CTIP2 to recruit both HDAC1/2 and the HP1-SUV39H1 complex^54^. Second, other factors directly compete with Sp1 for binding to the same GC-rich motifs including KLF2 and KLF3 in CD4⁺ T cells^55^, or Sp3 in myeloid cells^56^, which all recruit HDAC1/2 to enforce transcriptional silencing. Importantly, Sp3 is the only factor documented so far to directly compete with Sp1 for binding at these sites in myeloid cells. During myeloid differentiation, the Sp1/Sp3 ratio changes, from Sp3-dominant in monocytes to Sp1-dominant in macrophages, switching the HIV-1 promoter from a repressed to an active state^57^. Thus, the functional roles of the three Sp1 binding sites in controlling HIV-1 transcription are highly heterogeneous and context-dependent, varying according to cellular differentiation state and cell type.

Importantly, we here report that KLF16 exhibited several distinctive features that distinguish it from other known repressors acting through the Sp1 sites of the 5’LTR U3 region. Unlike c-Myc^53^, KLF2/3^43^, or Sp3^56^, which all recruit only HDAC1/2 to the 5’LTR, KLF16 couples histone deacetylation with H3K9 trimethylation by co-recruiting both the Sin3A-HDAC1 and the HP1α-SUV39H1 complex, establishing a more robust repressive chromatin environment. This context-independent activity across multiple cell types contrasts sharply with CTIP2, which has been shown to be restricted to microglia^5,54,58,59^ and distinguishes KLF16 from the cell type- or differentiation-dependent repressors previously identified. To our knowledge, KLF16 is the first factor shown to co-recruit both SUV39H1 and HP1α to the HIV-1 5’LTR in both T-lymphocytic and myeloid cells. Given that HP1β and HP1γ are also involved in HIV-1 transcriptional silencing^54,60^, future studies should investigate whether KLF16 similarly recruits these HP1 isoforms. Additionally, given the high expression of KLF16 in the brain^61^, its role in microglial HIV-1 reservoirs and its potential interplay with CTIP2 also warrant further investigation.

Among the three tandem Sp1 sites within the HIV-1 core promoter, Sp1 site III has emerged as a critical hub for both transcriptional activation (by recruiting the coactivator p300 and mediating the Sp1/NF-κB interaction^51,62^) and repression (serving as a recruitment platform for HDAC1^51^). Our results revealed that KLF16 exhibited preferential binding to Sp1 site III, thereby positioning KLF16 as a key regulator capable of directly competing with Sp1 binding to its functionally most critical site within the HIV-1 LTR. Although, additional Sp1 binding sites have been described within the HIV-1 provirus, including in the 5’LTR leader region^63^, in the *pol* gene *cis*-regulatory region^64^, and in the 3’LTR^12^, we showed that KLF16 exclusively targeted the 5’LTR core promoter region. These results highlight that KLF16 is highly selective for the HIV-1 enhancer-core promoter Sp1 sites, with a strong preference for the Sp1 site III, in contrast to the broader Sp1 occupancy profile across the provirus.

The identification of KLF16 reported here as a transcriptional repressor acting at the three tandem Sp1 sites in the HIV-1 5’LTR adds to the growing list of cellular factors that converge on these *cis*-elements, underscoring that they function as a critical regulatory hub controlling viral transcription. However, how the occupancy and function of these Sp1 sites are dynamically regulated across different stages of HIV-1 infection, from latency to productive replication, remains incompletely understood. While Sp1 is ubiquitously expressed, expression of other Sp1 site-binding factors is modulated by the state of cellular differentiation and activation. Indeed, Sp3 expression is enriched in monocytes and favours transcriptional repression^56^, whereas KLF2 and KLF3 expression is highest in quiescent T cells and decreases upon activation, correlating with their role in maintaining HIV-1 latency^43,55,65^. These dynamics support a model in which changes in the relative expression levels of Sp1 and its competing repressors determine whether the three 5’LTR Sp1 sites activate or suppress HIV-1 gene expression. Here, we showed that KLF16 expression displayed an unexpected activation-dependent pattern distinct from KLF2 and KLF3. More specifically, KLF16 expression was significantly reduced in memory CD4^+^ T cells from ART-treated PWH compared to PWOH, suggesting that KLF16 levels were shaped by the persistent immune activation and transcriptional reprogramming known to occur in CD4^+^ T cells from PWH despite suppressive ART. Following TCR triggering *in vitro*, KLF16 expression was increased, contrasting with the recently reported activation-induced downregulation of KLF2 and KLF3^43,55,65–67^. This different temporal pattern suggests that KLF16 functions in establishing or re-establishing HIV-1 latency during or following T-cell activation. Future work will elucidate the dynamic regulation of the Sp/KLF regulatory hub across the HIV-1 activation-latency spectrum.

In the present study, we further showed that retinoic acid reactivated HIV-1 from latency in myeloid cells in part by downregulating KLF16 expression. The intestinal mucosa is a critical site of HIV-1 persistence^68^, where RA produced by mucosal dendritic cells creates a permissive microenvironment for viral replication. While RA-mediated enhancement of HIV-1 replication has been attributed to modulation of CCR5 expression, mTOR-dependent post-entry signalling, and differential CD4⁺ T cells trafficking^47,69–71^, the transcriptional mechanisms by which RA sustains HIV-1 gene expression in gut-resident myeloid cells remained unclear. We here identified KLF16 downregulation as a novel mechanistic link between RA exposure and HIV-1 transcriptional reactivation in myeloid cells, associated with loss of KLF16-dependent repressive chromatin marks at the 5’LTR. Notably, this effect was myeloid cell-specific, as ATRA-treated T-lymphocytic cells maintained stable KLF16 expression, suggesting a lineage-restricted dependency. As complete elimination of RA from the intestinal microenvironment is biologically unfeasible, the molecular crosstalk between RA signalling and KLF16-mediated silencing represents a potential therapeutic avenue for reducing viral persistence in this anatomic reservoir.

Interestingly, in addition to be context-dependent^22,24,25,28,29,31,72^, KLF16-mediated transcriptional silencing exhibited striking specificity: while the LTRs from HIV-1, HERV-K and HTLV-1 all depend on Sp1 binding sites for transcriptional regulation^38,73,74^, we showed that KLF16 was only specifically recruited to and repressed the HIV-1 5’LTR. This specificity is therapeutically advantageous as HERV expression represents a significant source of chronic inflammation and immune dysregulation linked to cancer development^75–77^. To date, no selective KLF16 inhibitors have been identified. Developing such inhibitors would represent a promising avenue to achieve HIV reactivation without inadvertently unleashing potentially pathogenic HERV expression, thereby enabling more selective and safer latency-reversing strategies.

A limitation of our study is that KLF16 was identified through a DNA pull-down screen using nuclear extracts from cell lines rather than from PWH primary cells. This methodological approach may have introduced biases, as established cell lines do not fully recapitulate the transcriptional landscape of primary cells. Additionally, our present study focused on memory CD4^+^ T cells from PWH peripheral blood, as well as primary MdMs from PWOH. Since HIV-1 persistence and the expression of transcriptional regulators may differ between peripheral blood and tissue compartments such as the gut, it would be interesting in future studies to examine whether the KLF16-mediated regulation of HIV-1 transcription identified here extends to other anatomical HIV-1 reservoirs.

## Resource availability

## Lead contact

Further information and requests for resources and reagents should be directed to and will be fulfilled by the lead contact, Carine Van Lint.

## Materials availability

All materials used are available from the corresponding author on request.

## Data and code availability

The mass spectrometry raw data generated in this study have been deposited to the ProteomeXchange Consortium via the PRIDE^78^ partner repository with the dataset identifier PXD077517 and 10.6019/PXD077517. All R scripts used for mass spectrometry data filtering and JASPAR-based transcription factor binding site analysis are publicly available on GitHub (https://github.com/Maryam-Bendoumou/HIV1-LTR-pulldown-proteomics.git) and permanently archived on Zenodo under DOI: https://doi.org/10.5281/zenodo.19389205. Any additional information required to reanalyze the data reported in this paper is available from the lead contact upon request.

## Acknowledgments

C.V.L. acknowledges funding from the Belgian Fund for Scientific Research (FNRS, Belgium); the French INSERM agency “ANRS/Maladies infectieuses émergentes”; the *Université libre de Bruxelles* (ULB - Action de Recherche Concertée (ARC) grant); ViiV Healthcare; the BREACH Foundation; the Internationale Brachet Stiftung (IBS); The “Amis des Instituts Pasteur à Bruxelles”, asbl.; the Canadian Institutes of Health Research (CIHR project grant PTJ-195736); the US National Institutes of Health (NIH) (MDC grant 1UM1AI164562, co-funded by National Heart, Lung and Blood Institute, National Institute of Diabetes and Digestive and Kidney Diseases, National Institute of Neurological Disorders and Stroke, National Institute on Drug Abuse and the National Institute of Allergy and Infectious Diseases). C.V.L. is Research Director (“Directrice de recherches”) of the FNRS. The laboratory of C.V.L. is part of the ULB-Cancer Research Centre (U-CRC) (Faculty of Medicine, ULB). M.S. was funded by a doctoral fellowship from the Belgian “Fonds pour la formation à la Recherche dans l’Industrie et dans l’Agriculture (FRIA)” (FNRS) and next from the “Amis des Instituts Pasteur à Bruxelles”, asbl and from the “Fonds David et Alice Van Buuren” and “Fondation Jaumotte-Demoulin et de la Fondation Héger-Masson” foundations. M.B. was funded by a doctoral fellowship from the Belgian “Fonds pour la formation à la Recherche dans l’Industrie et dans l’Agriculture (FRIA)” and next from the “Amis des Instituts Pasteur à Bruxelles”, asbl and is currrently supported by a post-doctoral fellowship from a “PDR” grant from the FNRS. A.D. is supported by a post-doctoral fellowship from a “PDR” grant from the FNRS. LP is funded by a PhD fellowship from the ULB and TM is a “Aspirant” fellow of the FNRS. LN was supported by a “PDR” grant from the FNRS. This work was also supported by research funding to P.A. from the Canadian HIV Cure Enterprise Team Grant (CanCURE2.0 and 3.0) funded by the Canadian Institutes of Health Research (CIHR; #HB2-164064; BR4 197730); and CIHR project grants (PJT #153052; PJT #178127; PJ4-192194; PTJ-195736). Core facilities and PWH/PWOH cohorts were supported by the Fondation du CHUM and the Fonds de recherche Québec Santé (FRQS) HIV/AIDS and Infectious Diseases Network.

## Contributors

Conceived and designed the study: CVL. Conceived and designed the experiments: MS, MB, PA and CVL. Performed the experiments: MS, MB, AD, SK, EP, C-D N-Y, LP, CV, JD, TM, AF, MD and PR. Provided access to clinical samples and related information: J-PR. Wrote the manuscript: MB, AD, MS, PA and CVL. Supervised the study and acquired funding: PA and CVL. All authors read or provided comments on the manuscript. All data were validated by CVL, PA, MS, MB and AD.

## Data sharing statement

All the data/analyses presented in the current manuscript will be made available upon request to the corresponding author.

---

**Extended Data Figure 1.**
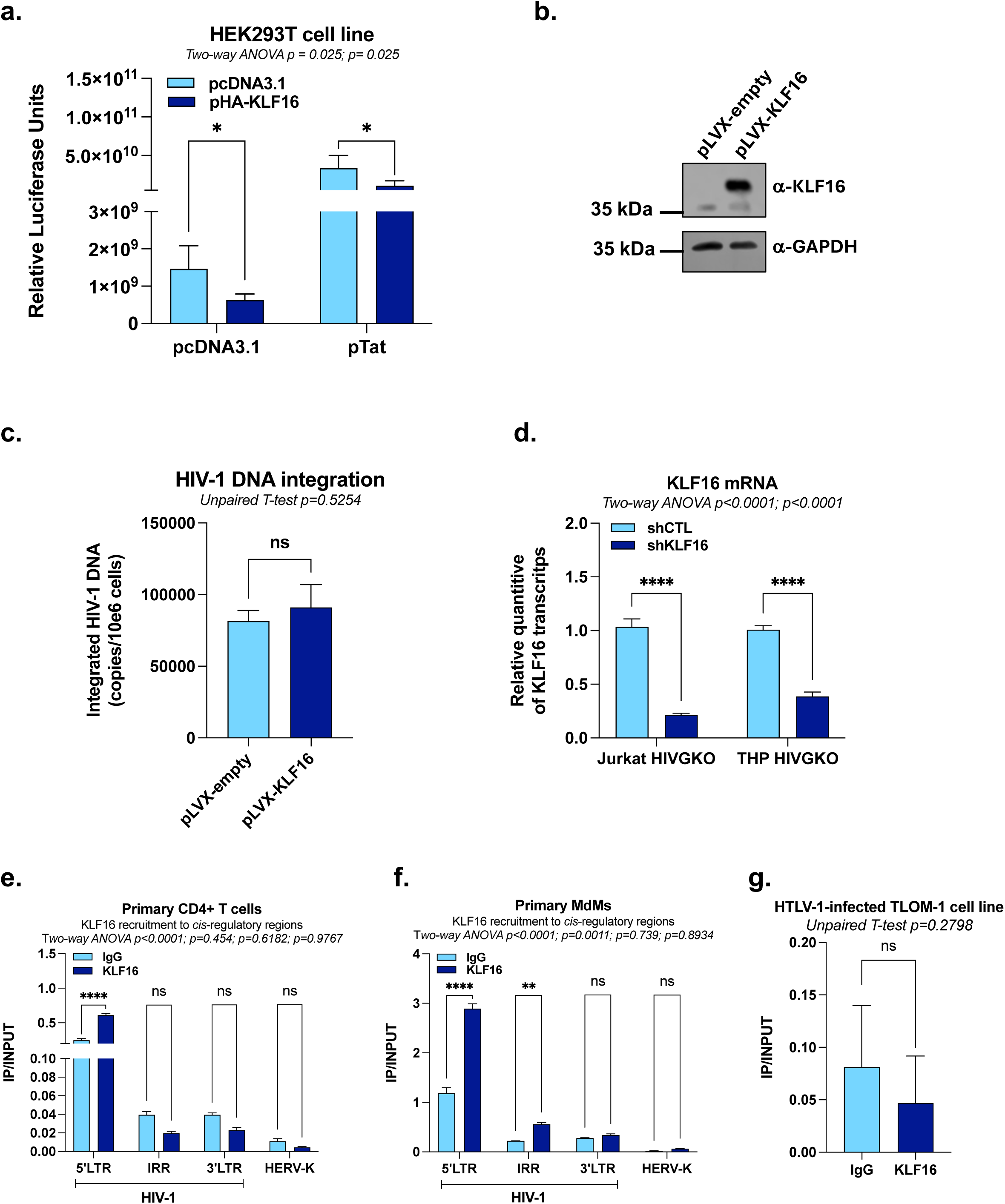
KLF16 is specifically recruited in vivo to the HIV-1 5’LTR and represses viral gene expression. **(a)** HEK293T cells were transiently co-transfected with pLTR-Fluc and either a single dose of pTat, a single dose of pHa-KLF16, or both. Luciferase activities were measured and normalized to protein concentration. **(b)** Verification of efficient KLF16 overexpression in the Jurkat cell lines pLVX-empty or pLVX-KLF16 by immunoblot using specific antibodies against KLF16 or GAPDH (serving as a loading control). **(c)** Uninfected Jurkat cell lines stably overexpressing either the control vector (pLVX-empty) or KLF16 (pLVX-KLF16) were infected with a fully replicative HIV-1 molecular clone, and the level of integrated HIV-1 provirus DNA was assessed by semi-nested RT-qPCR at day 2 post-infection. **(d)** Jurkat HIVGKO and THP HIVGKO cell lines transduced with either shCTL or shKLF16 were used for total RNA extraction to quantify KLF16 mRNA levels by RT-qPCR, normalized to TBP expression quantified at the same time. **(e)-(f)** Chromatin samples were prepared from primary CD4^+^ T cells or monocyte-derived-macrophages (MdMs) infected with HIV-1 and immunoprecipitated with a specific antibody against KLF16 or with a non-specific IgG serving as background control. KLF16 recruitment was assessed along the provirus using primers targeting the 5’LTR, the *pol* gene intragenic regulatory region (IRR) or the 3’LTR or assessed at the promoter of the endogenous retrovirus HERV-K. **(g)** Chromatin samples were prepared from the HTLV-1 latently-infected T-cell line TLOM-1 and were immunoprecipitated with a specific antibody against KLF16 or with a non-specific IgG serving as background control. KLF16 recruitment was assessed to each regulatory region using specific primers. One representative experiment out of three independent replicates is shown. Statistical analysis methods and corresponding *p* values are indicated above each graph.

**Extended Data Figure 2.**
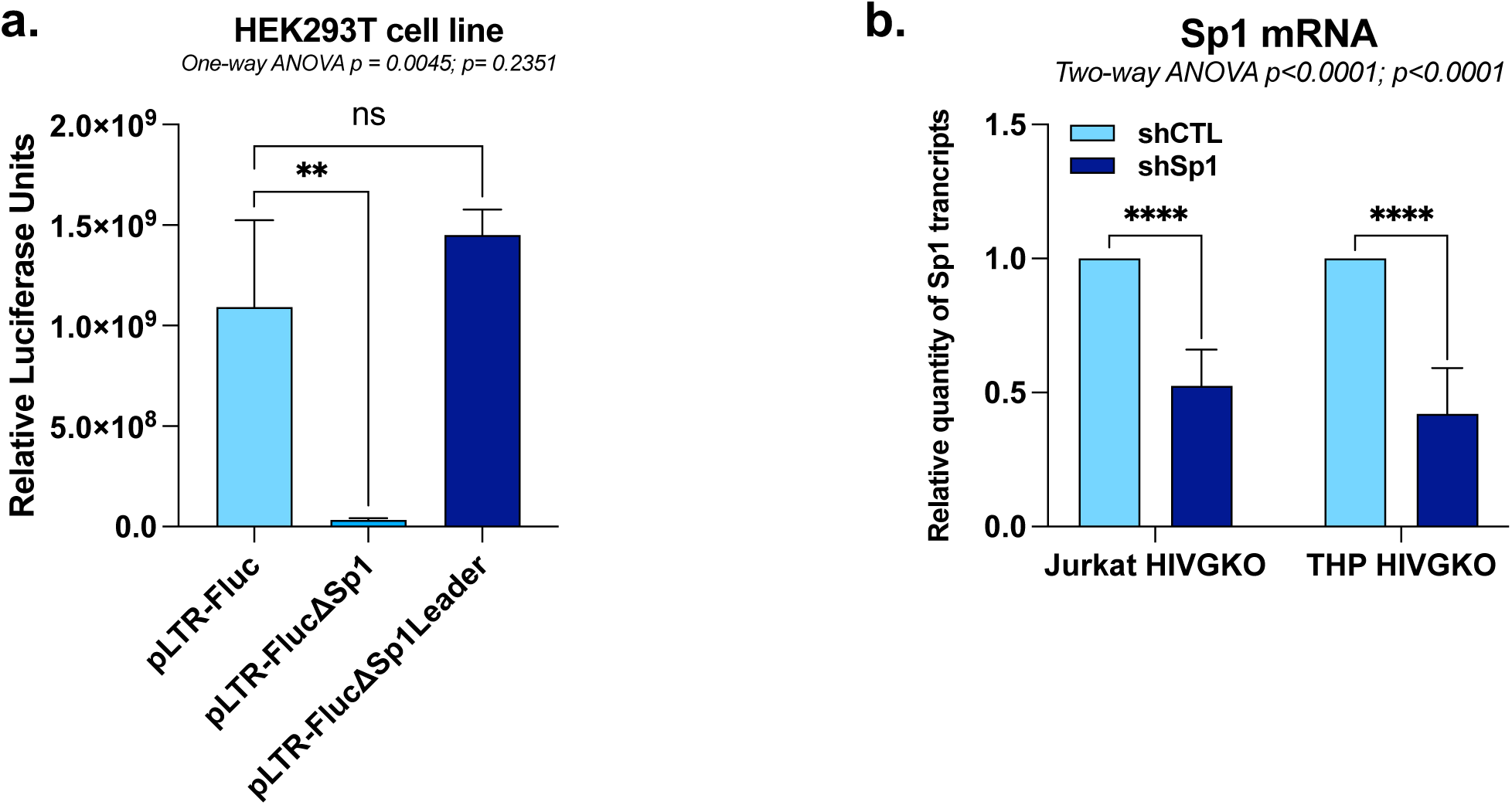
KLF16 competes with Sp1 for binding to the HIV-1 5’LTR to repress viral gene expression. **(a)** HEK293T cells were transiently transfected with either p-LTR-Fluc, either pLTR-FlucΔSp1, or pLTR-FlucΔSp1Leader. Luciferase activities were measured and normalized to protein concentration. **(b)** Jurkat HIVGKO and THP HIVGKO cells transduced with either shCTL or shSp1 were used for total RNA extraction to quantify KLF16 mRNA levels by RT-qPCR, normalized to TBP expression quantified at the same time. One representative experiment out of three independent replicates is shown. Statistical analysis methods and corresponding *p* values are indicated above each graph.

**Extended Data Figure 3.**
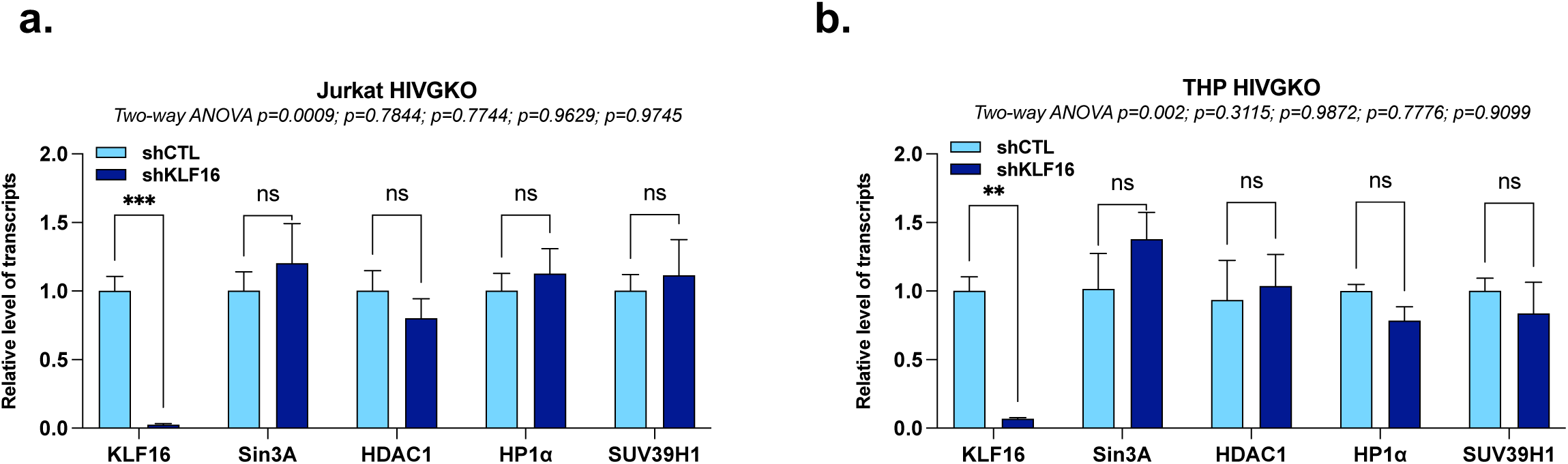
KLF16 modulates the epigenetic profile of the HIV-1 5’LTR to repress viral gene expression without impact on the expression of the epigenetic machineries. **(a)** and **(b)** Jurkat HIVGKO and THP HIVGKO cells transduced with either shCTL or shKLF16 were used for total RNA extraction to quantify KLF16, Sin3A, HDAC1, HP1α, and SUV39H1 mRNA levels by RT-qPCR, normalized to TBP expression quantified at the same time. One representative experiment out of three independent replicates is shown. Statistical analysis methods and corresponding *p* values are indicated above each graph.

## Materials and Methods

### Study participants

Leukapheresis samples were collected from PWOH and ART-treated PWH (**Supplementary Table S3**) at the McGill University Health Centre (MUHC) and Centre de recherche du Centre hospitalier de l’Université de Montréal (CR-CHUM, Montréal, Québec, Canada). PBMCs were isolated from leukapheresis by gradient centrifugation using the lymphocyte separation medium (Wisent, Saint-Jean-Baptiste/Canada). PBMCs were preserved frozen in liquid nitrogen in 10% DMSO (SIGMA, St. Louis/United States) and 90% fetal bovine serum (FBS; Wisent, Saint-Jean-Baptiste/Canada) until use.

### Ethics statement

Leukapheresis were collected from ART-treated PWH and PWOH, while respecting the principles included in the Declaration of Helsinki. This study was approved by the Institutional Review Board (IRB) of the McGill University Health Centre and the CHUM-Research Centre, Montreal, Quebec, Canada. The study participants provided a signed informed consents and agreed with the publication of results.

### Cell lines and cell culture

The T-lymphoid cell line Jurkat (RRID: CVCL_0065) and the monocytic cell line THP1 (RRID: CVCV_0006) were obtained from the AIDS Research and Reference Reagent Program (National Institute of Allergy and Infectious Diseases [NIAID], National Institutes of Health [NIH]). Using the Jurkat and THP1 cell lines, new HIV-1 latent cell lines (Jurkat HIVGKO and THP HIVGKO) were generated by single-round infection with the dual-reporter HIVGKO molecular clone, followed by flow-cytometric sorting of latently-infected cells (GFP⁻/mKO⁺), kindly provided by Eric Verdin^34^. Jurkat cells were also used to generate a Jurkat cell line overexpressing KLF16 (pLVX-KLF16) and the corresponding control cell line (pLVX-empty) by lentiviral transduction and puromycin selection of transduced cells. T-lymphocytic and monocytic cells were maintained in RPMI 1640-GlutaMAX medium (Life Technologies) supplemented with 10% fetal bovine serum (FBS). The human embryonic kidney cell line HEK293T (RRID: CVCL_0063) was obtained from the American Type Culture Collection (ATCC, Manassas, VA). The HEK293T cell line was maintained in Dulbecco’s modified Eagle’s-GlutaMAX medium (Life Technologies) supplemented with 10% FBS, with additional sodium pyruvate (1 mM). All media also contained 50 U/mL penicillin and 50 μg/mL streptomycin (Life Technologies). Cells were grown at 37°C in a humidified 95% air/5% CO₂ atmosphere.

### DNA-affinity approach and mass spectrometry analysis

The 100 bp desthiobiotinylated single-stranded sense and antisense oligonucleotides corresponding to a fragment of the HIV-1 5′ LTR (nt −102 to nt −2, where nt +1 is the start of the 5′ LTR U3 region) were produced by Integrated DNA Technologies (IDT) and subsequently annealed (Supplementary Table 2). DNA-affinity pull-down and trypsin digestion were conducted using a protocol previously reported by P. Renard and colleagues^79^. The digest was analyzed using nanoLC- MS/MS tims TOF Pro (Bruker, Billerica, MA, USA) coupled with an UHPLC nanoElute (Bruker). Peptides were separated by nanoLC (nanoElute, Bruker) on a 75 μm ID, 25 cm C18 column with integrated CaptiveSpray insert (Aurora, ionopticks, Melbourne) at a flow rate of 400 nl/min, at 50°C. Data dependent acquisition (DDA) on the tims TOF Pro was performed using Hystar 5.1 and timsControl 2.0. tims TOF Pro data were acquired using 160 ms TIMS accumulation time, mobility (1/K0) range from 0.7 to 1.4 Vs/cm². Mass-spectrometric analysis were carried out using the parallel accumulation serial fragmentation (PASEF) ^80^acquisition method. All MS/MS samples were analyzed using Mascot (Matrix Science, London, UK; version 2.7) and Scaffold (version Scaffold_4.10.0, Proteome Software Inc., Portland, OR) was used to validate MS/MS based peptide and protein identifications. Peptide identifications were accepted if they could be established at greater than 96.0% probability to achieve an FDR less than 1.0% by the Scaffold Local FDR algorithm.

### Antibodies and Drug

The antibodies and the drug (ATRA) used in this study are listed in **Supplementary Table 1**.

### Electrophoretic Mobility Shift Assays (EMSAs)

Nuclear extracts from Jurkat and THP-1 cells were prepared using a protocol described by Dignam and colleagues^81^ and protein concentrations were determined by Bradford assays (Bio-Rad, # 5000001). Pure extracts of KLF16 or Luciferase (used as a negative control) were prepared using the Transcription and Translation (TnT) system (Promega, #L1170). The oligonucleotide sequences used for the probes are shown in **Supplementary Table 2**. EMSAs, competition EMSAs and supershift assays were performed as described previously^82^.

### Plasmid constructs

The non-episomal reporter was generated by cloning the HIV-1 pNL4.3 LTR into the pGL2 luciferase reporter vector (Promega #E1611). The episomal HIV-1 5′ LTR (pLTR-Fluc) reporter plasmid was previously described by our laboratory^35^. The pLTR-FlucΔSp1 and pLTR-FlucΔSp1-leader vectors were generated by introducing mutations in Sp1 binding sites within the U3 and leader regions, as previously described^39,83^, *via* PCR fragments with mutated primers assembled using NEBuilder HiFi DNA Assembly Mix (NEB #E2621L). The pHA-KLF16 plasmid was generated by insertion of the KLF16 open reading frame (ORF) into the pcDNA3.1 expression vector using NEBuilder HiFi DNA Assembly Mix (NEB #E2621L). The pLVX-KLF16 and pLVX-empty vectors were generated from the pLVX-GFP control vector, kindly provided by Maud Martin (*Université libre de Bruxelles*, Brussels, Belgium), by excision of GFP using restriction enzymes, insertion of the KLF16 ORF from pKLF16, and assembly with NEBuilder HiFi DNA Assembly Mix (NEB #E2621L). Plasmid encoding for Sp1 (pSp1) was acquired from Addgene. The lentiviral shRNA plasmids (called shKLF16 and shCTL (non-targeting)) were obtained from VectorBuilder (VB900126-3570CSV and VB10000-0013dtn, respectively). TRC lentiviral shRNAs targeting Sp1 were obtained from Horizon Discovery (#RHS4533-EG6667). The envelope pVSG-G and packaging psPAX2 vectors were kindly provided by Prof. Angela Ciuffi (University of Lausanne, Lausanne, Switzerland). For all constructs, the cloned fragments were fully sequenced by Sanger sequencing.

### Transient transfection and luciferase assays

The Jurkat and THP1 cell lines were co-transfected with 200 ng of reporter construct and with increasing doses of an expression vector coding for the HA-KLF16 protein using the TurboFect transfection reagent (Thermofisher, #R0533). HEK293T cells were co-transfected with 200 ng of reporter vector and with 400 ng either of an expression plasmid coding for HA-KLF16 (pHA-KLF16) or of pcDNA3.1 (as empty control vector) and an increasing dose of pSp1 using the calcium phosphate transfection method according to the manufacturer’s protocol (Takara #631312). Forty-eight hours post-transfection, cells were harvested, lysed, and luciferase activities were measured using the SingleGlo luciferase reporter assay (Promega #E1500). Total protein concentrations were determined by Bradford assay (Bio-Rad #5000001). Results were normalized for transfection efficiency using total protein concentrations.

### RNA extraction and analysis of transcripts

Total RNA samples were isolated using the All-In-One DNA/RNA/Protein Miniprep Kit (Biobasic, #BS88003) according to the manufacturer’s protocol. Reverse transcription was performed with HiScript III RT SuperMix for qPCR following the manufacturer recommendation (Vazyme, #R323-01). cDNAs were quantified by quantitative PCR using Luna Universal qPCR Master Mix (NEB, # M3003S) in the QuantStudio 3 Real-Time PCR System (Applied Biosystems). The sequences of the primers used for quantification are listed in **Supplementary Table 2**.

### Lentiviral particles production and transduction assays

VSV-G pseudotyped particles (containing shCT, shKLF16, shSp1, pLVX-empty and pLVX-KLF16) were produced by transfection of HEK293T cells as described previously^35^. Jurkat and THP HIVGKO cells were transduced as described previously^84^. The selection of transduced cells was performed by completing the culture media with 1μg/mL of puromycin.

### Infection assays in cell lines

Stable Jurkat pLVX-empty and Jurkat pLVX-KLF16 cells were infected with the fully replicative HIV-1 strain 89.6, as described previously^85^. Following infection, cells were extensively washed and cultured in standard RPMI medium with 10% FBS; culture supernatants were harvested every two days for 10 days. HIV-1 replication kinetics were assessed by quantification of viral particles using SG-PERT assays, as described previously^36^. Integrated HIV-1 DNA was quantified as described previously^86^ and used to normalize HIV particle values, thereby correcting for initial infection variability between conditions.

### Chromatin immunoprecipitation assays (ChIP)

ChIP assays were performed as previously described^8^. Relative quantification using the standard curve method on the input was performed for each primer pair and 96-well Optical Reaction plates were read in a StepOnePlus PCR instrument (Applied Biosystems). Fold enrichments were calculated as fold inductions relative to the values measured with IgG. Primer sequences used for quantification are available upon request. The sequences of primers used for quantification are listed in **Supplementary Table 2**.

### Total protein extracts and Western Blotting

pLVX-empty and pLVX-KLF16 Jurkat cells were harvested and lysed in RIPA buffer completed with protease inhibitors (Roche). Total protein concentration was determined by *DC*™ Protein Assay Kit II (Biorad). Western blotting experiments were performed using 10 µg of total protein extracts. The immunodetections were assessed using primary antibodies targeting the proteins of interest. Horseradish peroxidase (HRP)-conjugated secondary antibodies were used for chemiluminescence detection (Cell Signaling Technology).

### Primary models for HIV-1 infection (study of KLF16 recruitment to promoters)

Buffy coats from HIV-uninfected donors were obtained from the *Centre de Transfusion du Sang de Charleroi* (Charleroi, Belgium). Peripheral blood mononuclear cells (PBMCs) were isolated by Ficoll gradient centrifugation (Stemcell, Lymphoprep #18061). CD4⁺ T cells were then isolated by negative magnetic bead selection (StemCell #19052), and CD14⁺ monocytes were isolated by positive magnetic bead selection (StemCell #17858), according to manufacturers’ instructions. Following isolation, CD4⁺ T cells were stimulated with anti-CD3 and anti-CD28 antibodies (T Cell TransAct™, Miltenyi Biotec, #130-111-160) for three days and then infected by spinoculation (2h, 800xg, 32°C) with VSV-G-pseudotyped HIVGKO^34^ particles at a multiplicity of infection (MOI) of 3000, where GFP and mKO2 expression were used to distinguish between productively-infected and latently-infected cells. After spinoculation, cells were allowed to recover for five days with conditioned medium replacement every 24 h to promote cell survival and quiescence. CD14⁺ monocytes were resuspended at 2×10⁶ cells/mL in RPMI supplemented with 20% fetal bovine serum (FBS), 10 ng/mL each of M-CSF and GM-CSF (ProteinTech #HZ-119 and #HZ-1002, respectively). Cells were incubated for 6 days at 37°C, with half-medium replacement daily. Seven days after plating, cells were infected with VSV-G-pseudotyped HIV-1 pNL4.3GFP molecular clone (NIH: ARP_11349) at an MOI of 3000, where GFP expression was used to identify productively-infected cells.

### Study of KLF16 expression in PWOH and PWH and the effect of KLF16 downregulation on HIV-1 expression in primary CD4+ T cells and the effect of ATRA on KLF16 expression in MdM

#### Memory CD4+ T cell enrichment and TCR activation

Memory CD4^+^ T cells were enriched from PBMCs of PWOH and ART-treated PWH by negative selection using magnetic beads (EasySep™ Human Memory CD4^+^ T Cell Enrichment Kit, #19157, STEMCELL Technologies, Vancouver, BC, Canada). Cell purity was analysed by flow cytometry upon staining with anti-CD3, anti-CD4 and anti-CD28 antibodies (**Supplementary Table S1**). Cells were activated using immobilized anti-CD3 and soluble anti-CD28 antibodies (1 µg/mL, BD Pharmingen, San Diego, CA, USA) for up to 3 days and used for subsequent experiments.

#### Monocyte-derived macrophage (MdMs) differentiation

Total monocytes were isolated from PBMCs of PWOH by negative selection using magnetic beads (Miltenyi), as we previously reported^47^. Cell purity was analysed by flow cytometry upon staining with CD3, CD4, HLA-DR, CD1c, CD14 and CD16 antibodies (**Supplementary Table S1**), with monocytes being identified as CD3^−^CD4^low^CD1c^−^HLA-DR^+^CD14^+/-^CD16^+/-^ cells. Monocytes were cultured in the presence of human recombinant M-CSF (20 ng/ml; R&D Systems) for 6 days in the presence or the absence of ATRA (**Supplementary Table S3**), with media being refreshed every 2 days, as we previously reported^47^. At day 6, macrophages acquired a typical fibroblastic morphology and a typical CD14^+^CD16^+^ phenotype, as we reported^47^.

#### Virus stocks production and quantification

Single-round infection was performed with vesicular stomatitis virus (VSV)-G-pseudotyped-HIV-1 (HIV_VSVG_) generated using two plasmids: ***i)*** the plasmid pHEF Expressing Vesicular Stomatitis Virus (VSV-G) (ARP-4693), obtained through the NIH HIV Reagent Program, Division of AIDS, NIAID, NIH, contributed by Dr. Lung-Ji Chang; and ***ii)*** the plasmid encoding for the HIV vector containing the NL4-3 backbone encoding for enhanced green fluorescent protein (EGFP) in place of the Envelope (Env) (NL4.3EGFPΔEnv), and encoding for functional Nef and Vpu, was obtained through the NIH HIV Reagent Program, Division of AIDS, NIAID, NIH, contributed by Dr. Haili Zhang, Dr. Yan Zhou and Dr. Robert Siliciano. HIV-1 stocks were generated by transfection in 293T using X-tremeGENE HP DNA Transfection Reagent (#6366236001, Roche Diagnostics, Mannheim, Germany) as we previously described^87^. Viral stock concentration was determined by ELISA quantification of soluble HIV-p24 levels in cell-culture supernatants, using a home-made ELISA, as we previously described^87^. Viral stock titration was performed by infection of TCR-activated primary CD4+ T cells and the % of infected cells was evaluated by flow cytometry upon intracellular staining with CD4 and HIV-p24 Abs, as we previously described^87^. The optimal viral titres were selected based on maximum % of productively infected cells (CD4_low_HIV-p24_+_) and minimum cell mortality evaluated using the viability dye AquaVivid.

#### KLF16 silencing

KLF16 siRNA interference was performed in primary CD4^+^ T cells, as we described previously^88^. Briefly, memory CD4^+^ T cells were stimulated by CD3/CD28 Abs for 2 days. Activated cells were nucleofected with 1.5 μg KLF16 or non-targeting (NT1) siRNA (ON-TARGETplus SMART pool, Dharmacon) using the Amaxa Human T cell Nucleofector Kit (Lonza, Walkersville, MD, USA), according to the manufacturer’s protocol. Nucleofected cells (2 × 10^6^) were cultured for 24 h at 37°C in the presence of IL-2 (5 ng/mL). Cells were exposed to HIV_VSV-G_ and cultured for 3 more days in the presence of IL-2. At day 3 post-infection, cells were stained with LIVE/DEAD Fixable Dead Cell Stain Kit (Vivid, Life Technologies,CA) were analyzed by FACS (BD LSRII) (**Supplementary Table S2: List of oligonucleotides used in this study**).

#### Real time RT-PCR quantification of KLF16 mRNA expression

The efficacy of KLF16 silencing was determined by SYBR Green real-time RT-PCR quantification of KLF16 mRNA expression, 24h post-nucleofection. For this, total RNA was isolated using the RNAeasy mini kit (Qiagen, Hilden, Germany). The KLF16 mRNA was quantified by one step SYBR Green real-time RT-PCR using the custom primers from IDT (Table KRT). The RT-PCR reactions were carried out using the Light Cycler 480 II (Roche, Basel, Switzerland). The relative expression of KLF16 was normalized to the housekeeping gene 28S rRNA (**Supplementary Table S2: List of oligonucleotides used in this study)**.

#### Integrated HIV-DNA quantification

Integrated HIV-DNA levels were quantified by nested real-time PCR (Light Cycler 480, Roche) (**Supplementary Table S2: List of oligonucleotides used in this study**). Experiments were performed in triplicates. HIV-1 copy numbers were normalized to CD3 copy numbers (two CD3 copies per cell), with the limit of detection being 3 HIV-DNA copies/reaction, as we previously described^87^.

#### Flow cytometry analysis

For flow cytometry analysis, antibodies (Abs) were listed in the Key Resource Table. Intracellular staining was performed with BD Cytofix/Cytoperm kit (BD Biosciences, Franklin Lakes, NJ, USA), as we previously described (PMID: 39252967). The viability dye LIVE/DEAD Fixable Aqua Dead Cell Stain Kit (Vivid, Life Technologies, CA, USA) was used to exclude dead cells. Flow cytometry analysis was performed using an LSRII cytometer (BD Pharmingen San Diego, CA, USA). The positivity gates were placed using the fluorescence minus one (FMO) strategy. All flow cytometry data were analyzed with the FlowJo software version 10.10.0 (Tree Star, Inc).

---

**Supplementary Table S1:**
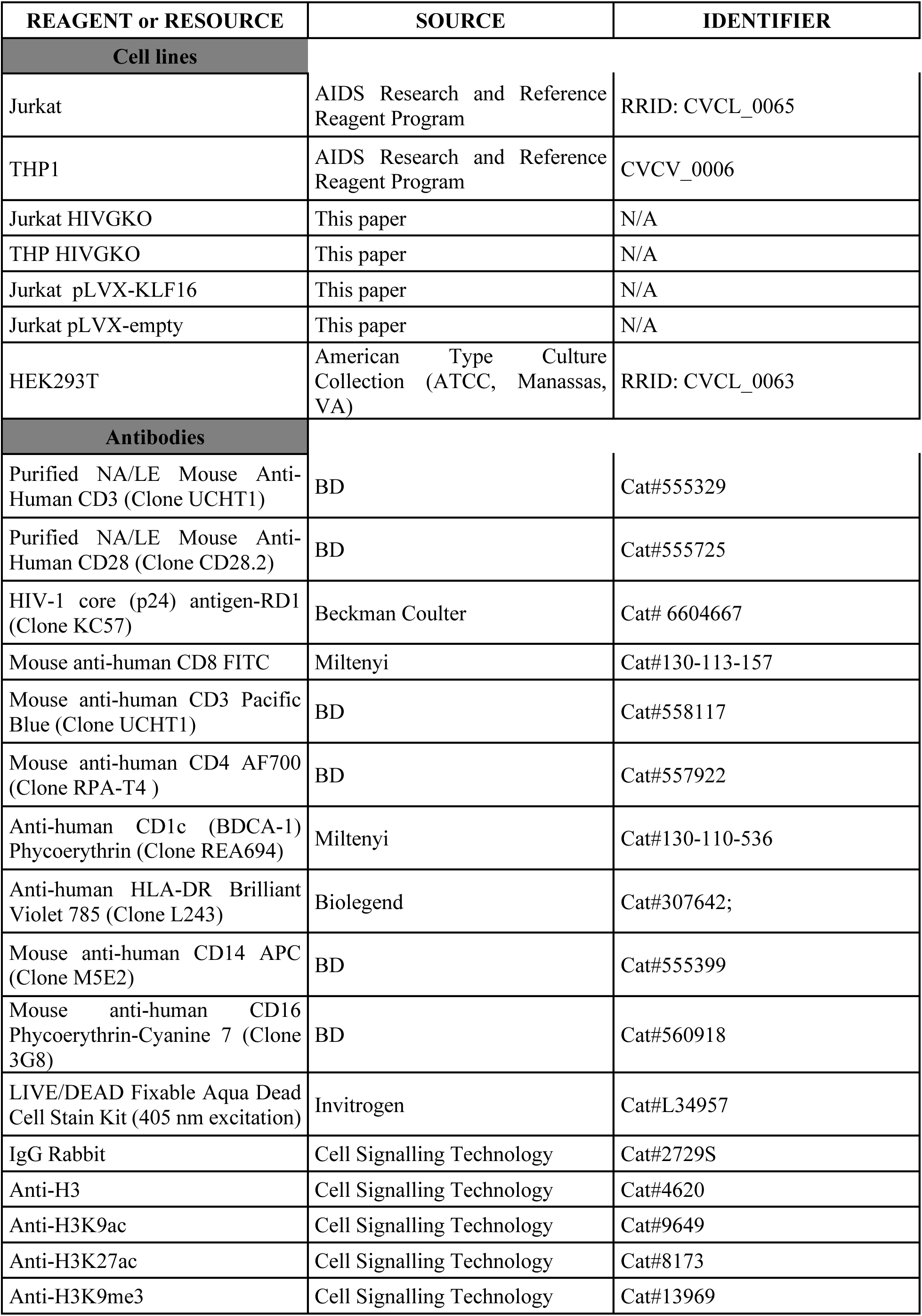
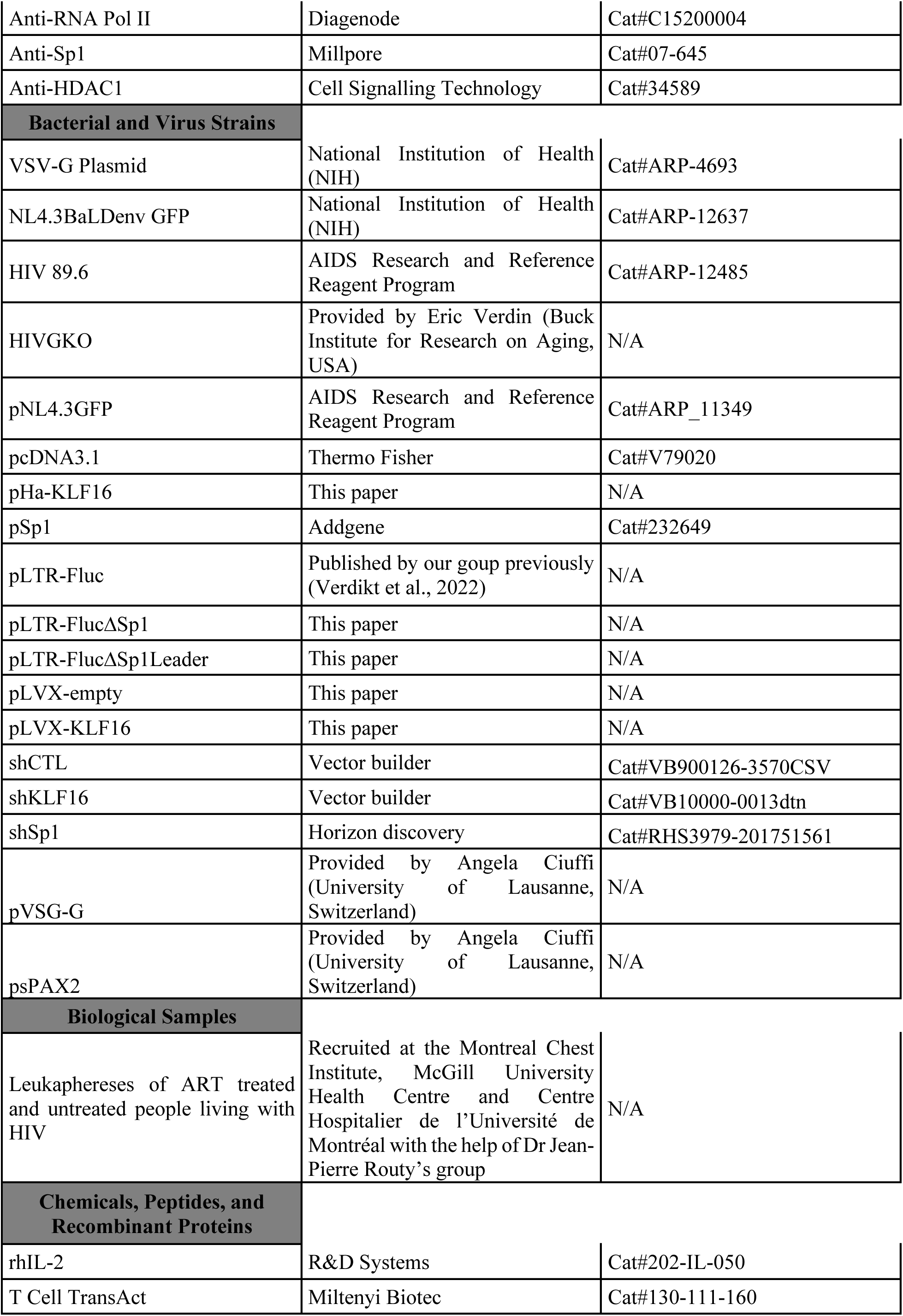
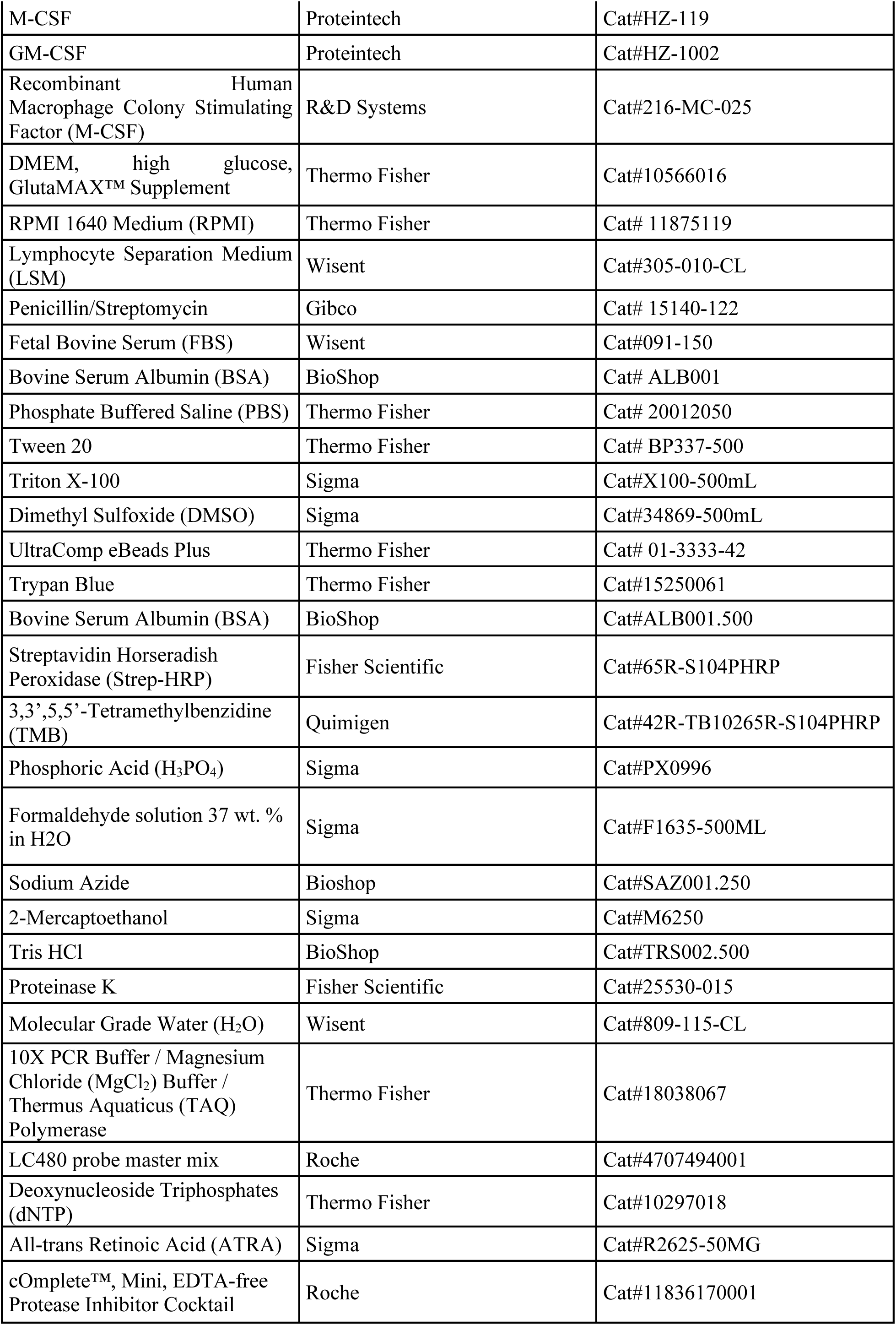
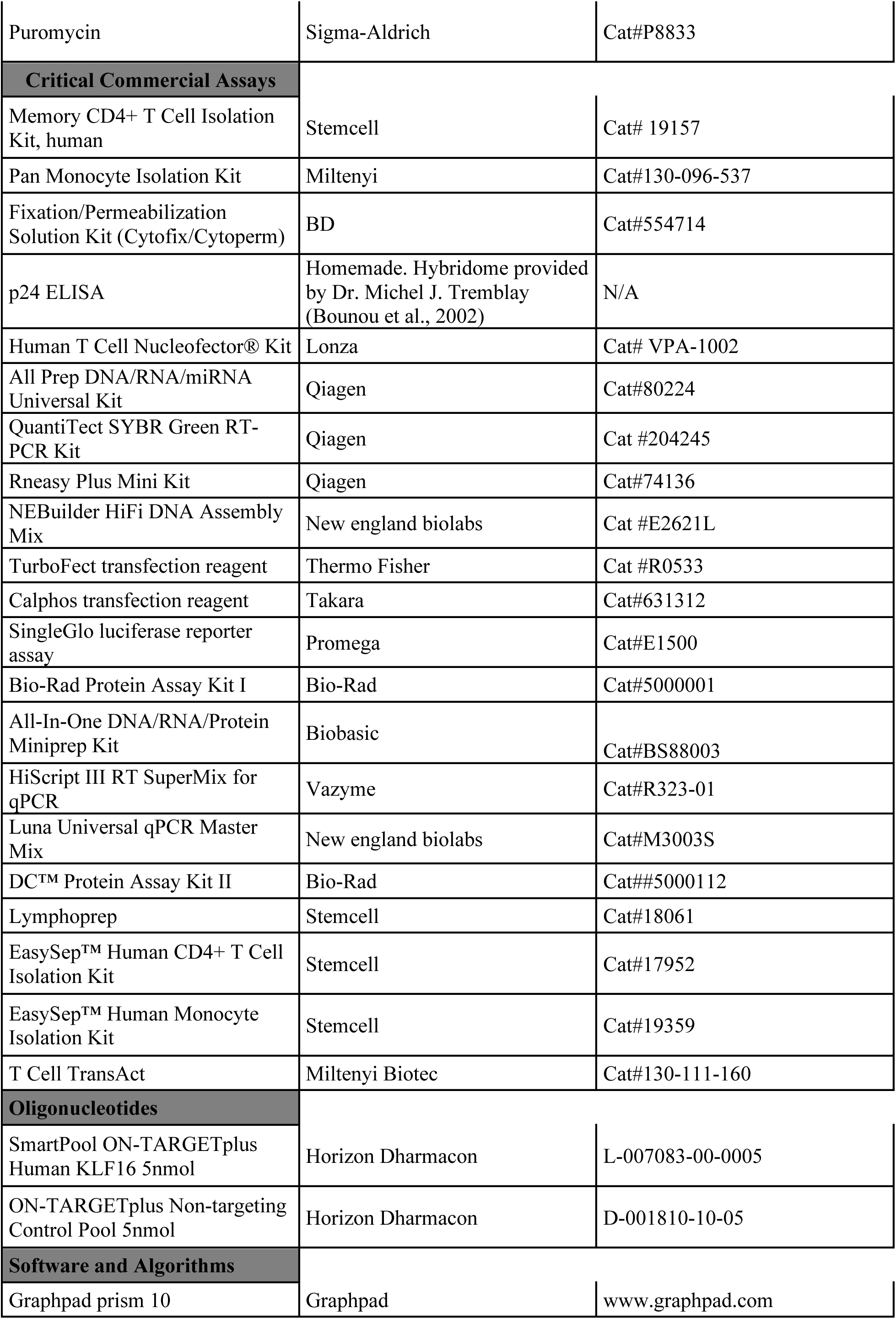
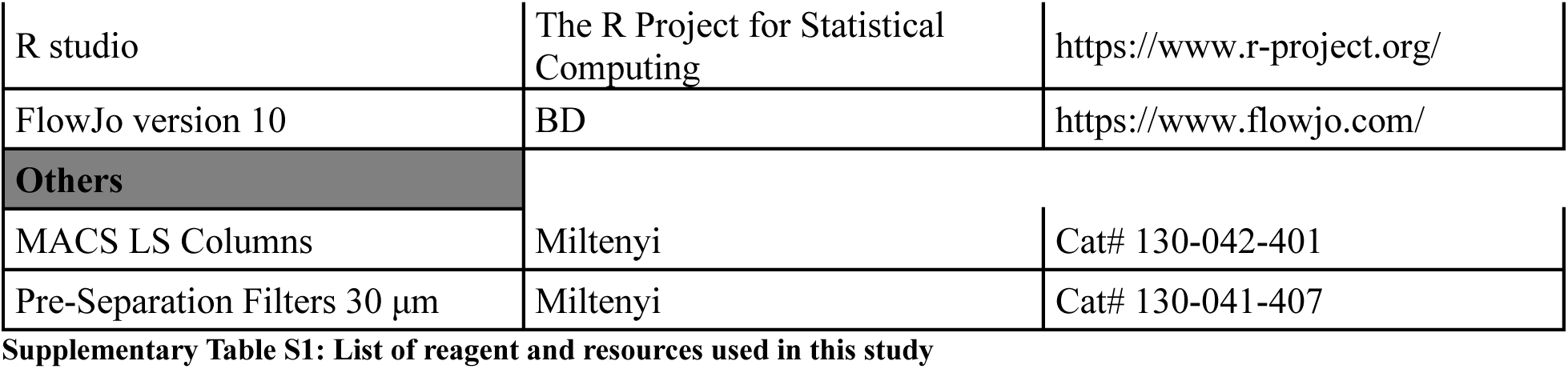
List of reagent and resources used in this study.

| REAGENT or RESOURCE | SOURCE | IDENTIFIER |
| --- | --- | --- |
| **Cell lines** |  |  |
| Jurkat | AIDS Research and Reference Reagent Program | RRID: CVCL_0065 |
| THP1 | AIDS Research and Reference Reagent Program | CVCV_0006 |
| Jurkat HIVGKO | This paper | N/A |
| THP HIVGKO | This paper | N/A |
| Jurkat pLVX-KLF16 | This paper | N/A |
| Jurkat pLVX-empty | This paper | N/A |
| HEK293T | American Type Culture Collection (ATCC, Manassas, VA) | RRID: CVCL_0063 |
| **Antibodies** |  |  |
| Purified NA/LE Mouse Anti-Human CD3 (Clone UCHT1) | BD | Cat#555329 |
| Purified NA/LE Mouse Anti-Human CD28 (Clone CD28.2) | BD | Cat#555725 |
| HIV-1 core (p24) antigen-RD1 (Clone KC57) | Beckman Coulter | Cat# 6604667 |
| Mouse anti-human CD8 FITC | Miltenyi | Cat#130-113-157 |
| Mouse anti-human CD3 Pacific Blue (Clone UCHT1) | BD | Cat#558117 |
| Mouse anti-human CD4 AF700 (Clone RPA-T4 ) | BD | Cat#557922 |
| Anti-human CD1c (BDCA-1) Phycoerythrin (Clone REA694) | Miltenyi | Cat#130-110-536 |
| Anti-human HLA-DR Brilliant Violet 785 (Clone L243) | Biolegend | Cat#307642; |
| Mouse anti-human CD14 APC (Clone M5E2) | BD | Cat#555399 |
| Mouse anti-human CD16 Phycoerythrin-Cyanine 7 (Clone 3G8) | BD | Cat#560918 |
| LIVE/DEAD Fixable Aqua Dead Cell Stain Kit (405 nm excitation) | Invitrogen | Cat#L34957 |
| IgG Rabbit | Cell Signalling Technology | Cat#2729S |
| Anti-H3 | Cell Signalling Technology | Cat#4620 |
| Anti-H3K9ac | Cell Signalling Technology | Cat#9649 |
| Anti-H3K27ac | Cell Signalling Technology | Cat#8173 |
| Anti-H3K9me3 | Cell Signalling Technology | Cat#13969 |
| Anti-RNA Pol II | Diagenode | Cat#C15200004 |
| Anti-Sp1 | Millipore | Cat#07-645 |
| Anti-HDAC1 | Cell Signalling Technology | Cat#34589 |
| **Bacterial and Virus Strains** |  |  |
| VSV-G Plasmid | National Institution of Health (NIH) | Cat#ARP-4693 |
| NL4.3BaLDenv GFP | National Institution of Health (NIH) | Cat#ARP-12637 |
| HIV 89.6 | AIDS Research and Reference Reagent Program | Cat#ARP-12485 |
| HIVGKO | Provided by Eric Verdin (Buck Institute for Research on Aging, USA) | N/A |
| pNL4.3GFP | AIDS Research and Reference Reagent Program | Cat#ARP_11349 |
| pcDNA3.1 | Thermo Fisher | Cat#V79020 |
| pHa-KLF16 | This paper | N/A |
| pSp1 | Addgene | Cat#232649 |
| pLTR-Fluc | Published by our group previously (Verdikt et al., 2022) | N/A |
| pLTR-FlucΔSp1 | This paper | N/A |
| pLTR-FlucΔSp1Leader | This paper | N/A |
| pLVX-empty | This paper | N/A |
| pLVX-KLF16 | This paper | N/A |
| shCTL | Vector builder | Cat#VB900126-3570CSV |
| shKLF16 | Vector builder | Cat#VB10000-0013dtn |
| shSp1 | Horizon discovery | Cat#RHS3979-201751561 |
| pVSG-G | Provided by Angela Ciuffi (University of Lausanne, Switzerland) | N/A |
| psPAX2 | Provided by Angela Ciuffi (University of Lausanne, Switzerland) | N/A |
| **Biological Samples** |  |  |
| Leukaphereses of ART treated and untreated people living with HIV | Recruited at the Montreal Chest Institute, McGill University Health Centre and Centre Hospitalier de l'Université de Montréal with the help of Dr Jean-Pierre Routy's group | N/A |
| **Chemicals, Peptides, and Recombinant Proteins** |  |  |
| rhIL-2 | R&D Systems | Cat#202-IL-050 |
| T Cell TransAct | Miltenyi Biotec | Cat#130-111-160 |
| M-CSF | Proteintech | Cat#HZ-119 |
| GM-CSF | Proteintech | Cat#HZ-1002 |
| Recombinant Human Macrophage Colony Stimulating Factor (M-CSF) | R&D Systems | Cat#216-MC-025 |
| DMEM, high glucose, GlutaMAX™ Supplement | Thermo Fisher | Cat#10566016 |
| RPMI 1640 Medium (RPMI) | Thermo Fisher | Cat# 11875119 |
| Lymphocyte Separation Medium (LSM) | Wisent | Cat#305-010-CL |
| Penicillin/Streptomycin | Gibco | Cat# 15140-122 |
| Fetal Bovine Serum (FBS) | Wisent | Cat#091-150 |
| Bovine Serum Albumin (BSA) | BioShop | Cat# ALB001 |
| Phosphate Buffered Saline (PBS) | Thermo Fisher | Cat# 20012050 |
| Tween 20 | Thermo Fisher | Cat# BP337-500 |
| Triton X-100 | Sigma | Cat#X100-500mL |
| Dimethyl Sulfoxide (DMSO) | Sigma | Cat#34869-500mL |
| UltraComp eBeads Plus | Thermo Fisher | Cat# 01-3333-42 |
| Trypan Blue | Thermo Fisher | Cat#15250061 |
| Bovine Serum Albumin (BSA) | BioShop | Cat#ALB001.500 |
| Streptavidin Horseradish Peroxidase (Strep-HRP) | Fisher Scientific | Cat#65R-S104PHRP |
| 3,3',5,5'-Tetramethylbenzidine (TMB) | Quimigen | Cat#42R-TB10265R-S104PHRP |
| Phosphoric Acid (H<sub>3</sub>PO<sub>4</sub>) | Sigma | Cat#PX0996 |
| Formaldehyde solution 37 wt. % in H<sub>2</sub>O | Sigma | Cat#F1635-500ML |
| Sodium Azide | Bioshop | Cat#SAZ001.250 |
| 2-Mercaptoethanol | Sigma | Cat#M6250 |
| Tris HCl | BioShop | Cat#TRS002.500 |
| Proteinase K | Fisher Scientific | Cat#25530-015 |
| Molecular Grade Water (H<sub>2</sub>O) | Wisent | Cat#809-115-CL |
| 10X PCR Buffer / Magnesium Chloride (MgCl<sub>2</sub>) Buffer / Thermus Aquaticus (TAQ) Polymerase | Thermo Fisher | Cat#18038067 |
| LC480 probe master mix | Roche | Cat#4707494001 |
| Deoxynucleoside Triphosphates (dNTP) | Thermo Fisher | Cat#10297018 |
| All-trans Retinoic Acid (ATRA) | Sigma | Cat#R2625-50MG |
| cOmplete™, Mini, EDTA-free Protease Inhibitor Cocktail | Roche | Cat#11836170001 |
| Puromycin | Sigma-Aldrich | Cat#P8833 |
| **Critical Commercial Assays** |  |  |
| Memory CD4+ T Cell Isolation Kit, human | Stemcell | Cat# 19157 |
| Pan Monocyte Isolation Kit | Miltenyi | Cat#130-096-537 |
| Fixation/Permeabilization Solution Kit (Cytofix/Cytoperm) | BD | Cat#554714 |
| p24 ELISA | Homemade. Hybridome provided by Dr. Michel J. Tremblay (Bounou et al., 2002) | N/A |
| Human T Cell Nucleofector® Kit | Lonza | Cat# VPA-1002 |
| All Prep DNA/RNA/miRNA Universal Kit | Qiagen | Cat#80224 |
| QuantiTect SYBR Green RT-PCR Kit | Qiagen | Cat #204245 |
| Rneasy Plus Mini Kit | Qiagen | Cat#74136 |
| NEBuilder HiFi DNA Assembly Mix | New england biolabs | Cat #E2621L |
| TurboFect transfection reagent | Thermo Fisher | Cat #R0533 |
| Calphos transfection reagent | Takara | Cat#631312 |
| SingleGlo luciferase reporter assay | Promega | Cat#E1500 |
| Bio-Rad Protein Assay Kit I | Bio-Rad | Cat#5000001 |
| All-In-One DNA/RNA/Protein Miniprep Kit | Biobasic | Cat#BS88003 |
| HiScript III RT SuperMix for qPCR | Vazyme | Cat#R323-01 |
| Luna Universal qPCR Master Mix | New england biolabs | Cat#M3003S |
| DC™ Protein Assay Kit II | Bio-Rad | Cat##5000112 |
| Lymphoprep | Stemcell | Cat#18061 |
| EasySep™ Human CD4+ T Cell Isolation Kit | Stemcell | Cat#17952 |
| EasySep™ Human Monocyte Isolation Kit | Stemcell | Cat#19359 |
| T Cell TransAct | Miltenyi Biotec | Cat#130-111-160 |
| **Oligonucleotides** |  |  |
| SmartPool ON-TARGETplus Human KLF16 5nmol | Horizon Dharmacon | L-007083-00-0005 |
| ON-TARGETplus Non-targeting Control Pool 5nmol | Horizon Dharmacon | D-001810-10-05 |
| **Software and Algorithms** |  |  |
| Graphpad prism 10 | Graphpad | www.graphpad.com |
| R studio | The R Project for Statistical Computing | https://www.r-project.org/ |
| FlowJo version 10 | BD | https://www.flowjo.com/ |
| **Others** |  |  |
| MACS LS Columns | Miltenyi | Cat# 130-042-401 |
| Pre-Separation Filters 30 µm | Miltenyi | Cat# 130-041-407 |

**Supplementary Table S2:**
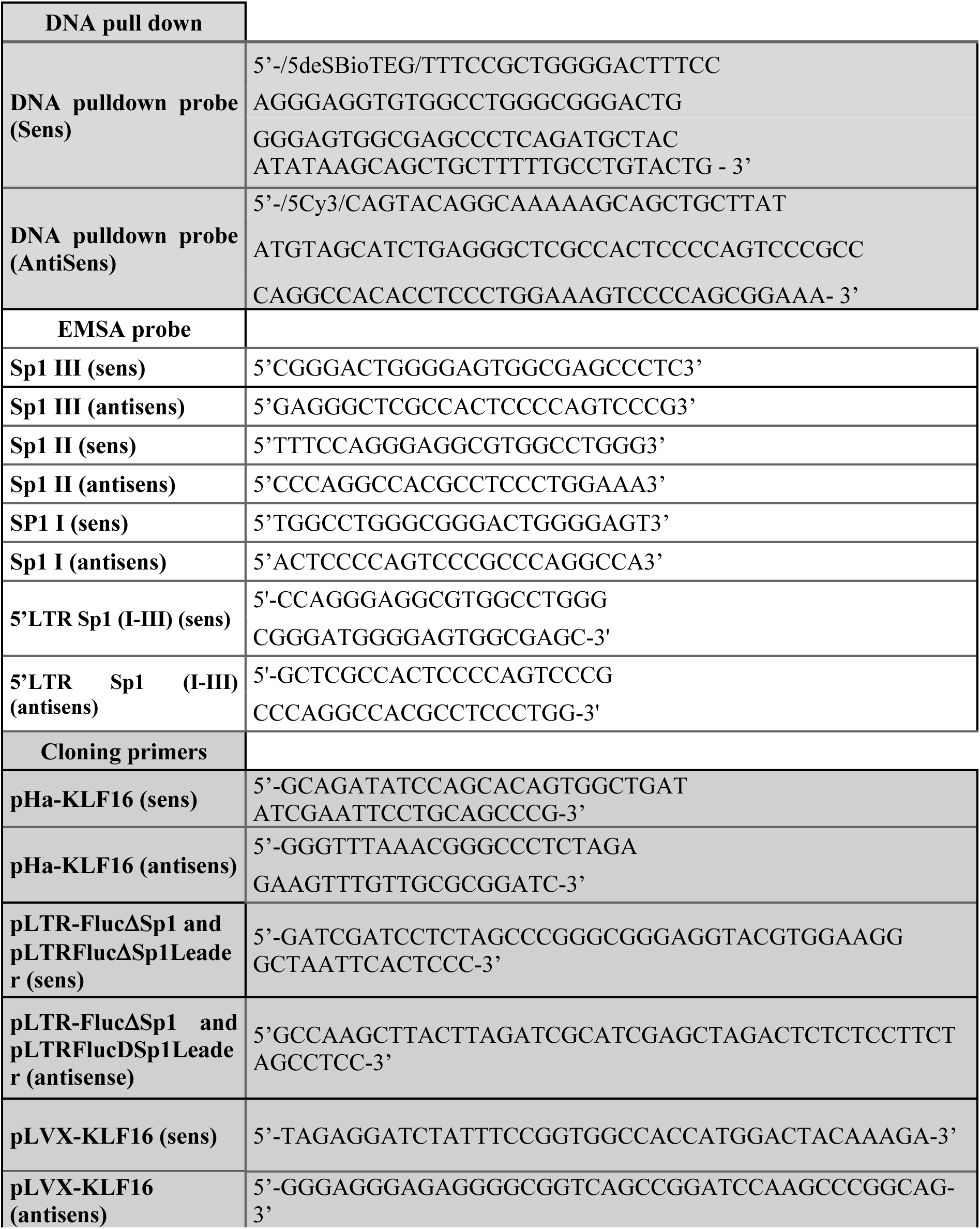
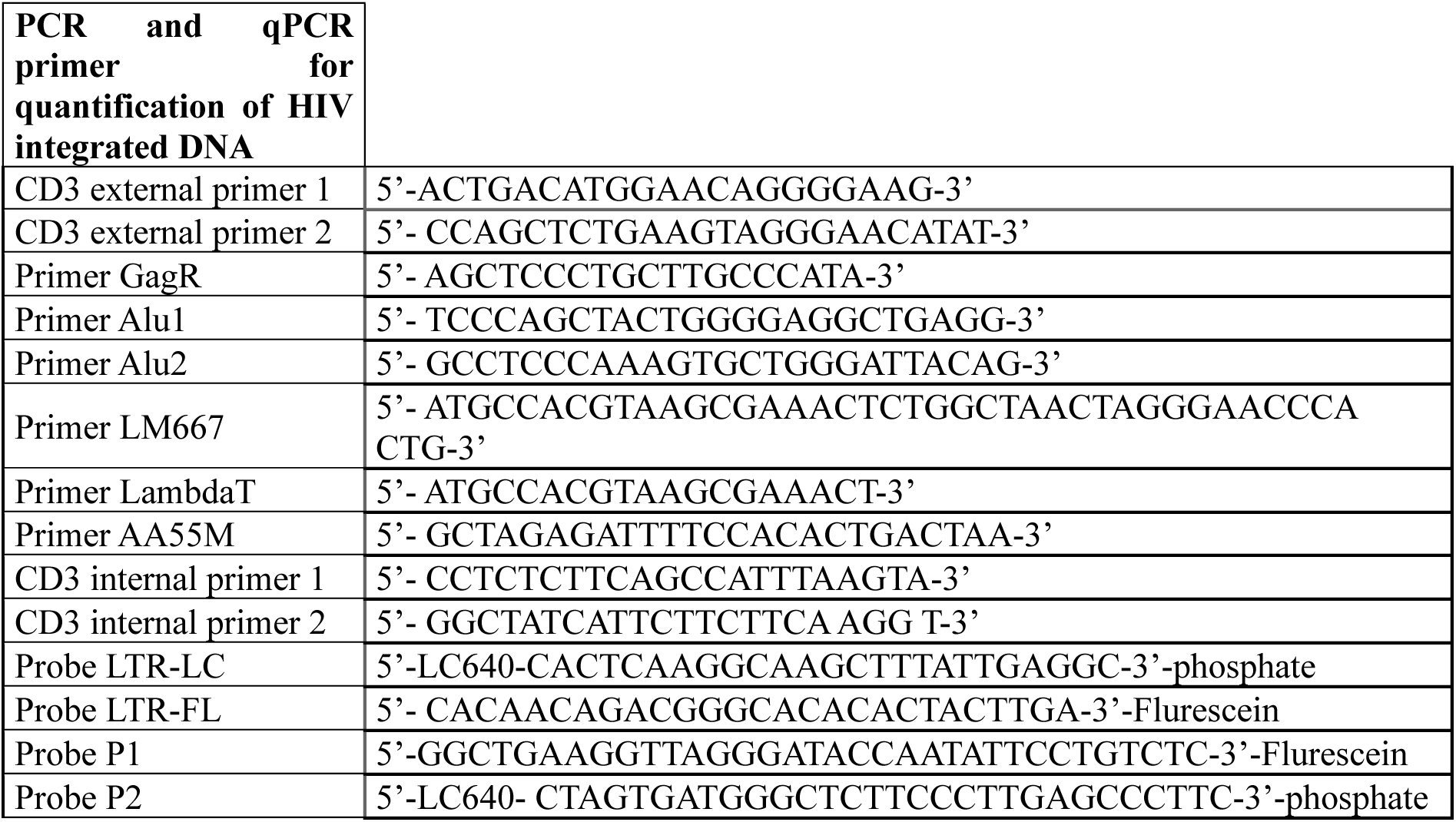
List of oligonucleotides used in this study.

**Supplementary Table S3:** Clinical parameters of HIV-infected study participants receiving viral suppressive antiretroviral therapy (ART).

| Participant<br>ID | Sex | CD4<br>counts<sup>#</sup> | CD8<br>counts<sup>#</sup> | Plasma<br>viral<br>load<sup>&</sup> | Time since<br>infection<sup>*</sup> | ART |
| --- | --- | --- | --- | --- | --- | --- |
| Donor #1 | M | 398 | 775 | <40 | 154 | Complera |
| Donor #2 | M | 542 | 803 | <40 | 13 | Stribild |
| Donor #3 | M | 743 | 899 | <40 | 171 | Truvada/Raltegravir |
| Donor #4 | M | 841 | 1322 | <40 | 149 | Sustiva/Truvada |
| Donor #5 | M | 908 | 854 | <40 | 89 | Stribild |
| Donor #6 | M | 963 | 644 | <40 | 123 | Prevista/Kivexa/Norvir |
| Donor #7 | M | 288 | 407 | <40 | 211 | Atripla |
| Donor #8 | M | 649 | 620 | <40 | 186 | Triumeq |

<sup>#</sup>, cells/ $\mu$ l; <sup>&</sup>, HIV RNA copies per ml plasma; <sup>\*</sup>, months; ART, antiretroviral therapy; <sup>§</sup>, months; NA,
information not available; ND, not detected

